# Characterization and evolutionary history of novel SARS-CoV-2-related viruses in bats from Cambodia

**DOI:** 10.1101/2025.04.15.648942

**Authors:** Tey Putita Ou, Julia Guillebaud, Artem Baidaliuk, Sothyra Tum, Dany Chheang, Deborah Delaune, Matthieu Prot, Rafael Rahal Guaragna Machado, Leakhena Pum, Vibol Hul, Thavry Hoem, Sovann Ly, Heidi Auerswald, Gavin James Smith, Philippe Dussart, Erik A Karlsson, Véronique Chevalier, Julien Cappelle, Veasna Duong, Etienne Simon-Lorière

**Affiliations:** Virology Unit, Institut Pasteur du Cambodge, Pasteur Network, Phnom Penh, Cambodia; Institut Pasteur, Université Paris Cité, CNRS UMR2000, Evolutionary Genomics of RNA Viruses unit, Paris, France; Institut Pasteur, Université Paris Cité, Bioinformatics and Biostatistics Hub, Paris, France; General Directorate of Animal Health and Production, Ministry of Agriculture, Forestry and Fisheries, Cambodia; Forestry Administration, Ministry of Agriculture, Forestry and Fisheries, Cambodia; Communicable Diseases Control Department, Ministry of Health, Cambodia; Duke-NUS Medical School, Programme in Emerging Infectious Diseases, Singapore; Institut Pasteur de Madagascar, Pasteur Network, Antananarivo, Madagascar; ASTRE (Animal, Santé, Territoires, Risques, Ecosystèmes), CIRAD, INRAE, Université de Montpellier, Montpellier, France; ASTRE (Animal, Santé, Territoires, Risques, Ecosystèmes), Antananarivo, Madagascar

## Abstract

Circulating bat coronaviruses present a significant pandemic threat, yet our understanding of their genetic diversity and evolutionary dynamics remains limited. Over 3 years, we sampled 1,462 bats in Cambodia’s Steung Treng province, identifying extensive and diverse coronaviruses co-circulation. Using metatranscriptomic and amplicon sequencing, we generated 33 complete sarbecovirus genomes, revealing novel lineages that cluster into four distinct groups, each associated with different *Rhinolophus* bat species. Our analysis highlights rapid migration and recombination of sarbecovirus lineages over short distances and timescales. Of note, the receptor-binding domains of two novel viral groups exhibit high similarity to SARS-CoV-2, and pseudovirus assays confirmed the ability of this spike protein to mediate entry into cells expressing human ACE2, suggesting a potential zoonotic risk. The observed genetic diversity underscores the urgent need for continuous surveillance to identify high-risk animal-to-human interfaces and inform pandemic preparedness.

## Introduction

The emergence of SARS-CoV and SARS-CoV-2 in recent decades has significantly underscored the zoonotic potential inherent within coronaviruses, particularly those of the sarbecovirus subgenus^1^. These events substantiated the urgent necessity to explore the extensive genetic diversity of coronaviruses, especially within wildlife reservoirs, to better understand the mechanisms driving their evolution and transmission. A comprehensive understanding of the diversity and distribution of sarbecoviruses is essential to assess their potential spillover risks and the mechanisms by which they might cross regions and species boundaries, and to inform public health strategies aimed at preventing future pandemics^1^.

Since the onset of the COVID-19 pandemic, our knowledge of sarbecoviruses has expanded, including the detection of these viruses in various animal species^2,3^ and notably among insectivorous horseshoe bats (genus *Rhinolophus*) across Asia^4^. Close relatives of SARS-CoV-2 have been identified in several regions, including China (in *R. affinis, R. stheno, R. sinicus, R. marshalli, R. malayanus*, and *R. pusillus)*^5–11^, but also in Japan (*R. cornutus*)^12^, in Thailand (*R. acuminatus*)^13^, in Cambodia (*R. shameli*)^14^, in Laos (*R. malayanus, R. marshalli*, and *R. pusillus*)^15^ and in Vietnam (*R. pusillus* and *R. affinis*)^16^. Some of these *Rhinolophus* bats have relatively wide geographical ranges and are known to co-roost in large numbers^11^. This behavior may facilitate viral transmission and recombination, highlighting the need for further research on their ecology. In the biodiverse regions of Southeast Asia, bats play important ecological roles (e.g. pollination, seed dispersal, pest control), and represent one of the most abundant groups of mammals^17^. Their role as reservoirs of viruses including coronaviruses^4^, and how their ecology influence the spread, maintenance, and evolution of these potentially zoonotic viruses remain insufficiently studied. The complexity of ecological interfaces in these regions raises concerns about the potential transmission of coronaviruses from their bat reservoir to other mammalian hosts and ultimately the risk of future sarbecovirus emergence in humans^18^. Effective monitoring of coronavirus circulation in bat populations before their emergence in humans or other animals, alongside understanding the molecular and epidemiological factors influencing spillover events^11^, remains a critical public health challenge.

In this study, we conducted longitudinal monitoring of coronaviruses in bats from Cambodia’s Steung Treng province. This region is characterized by karst hills formations where *Rhinolophus* spp. populations are known to roost, and where positive samples were previously identified in 2010^14^. Our study aimed to characterize the sarbecoviruses circulating in the region and investigate their evolutionary history and their potential zoonotic risk.

## Results

### Sample collection and testing

Between August 2020 and June 2023, we conducted eight field missions and captured and sampled 1,462 bats from nine genera (Extended Data Table 1). Of the 1,462 rectal swab samples tested, 146 were positive with at least one method: 23 tested positive exclusively by sarbecoviruses RT-qPCR^19^, 56 tested positive for pan-coronaviruses by at least one of two RT-PCRs^20,21^, and 67 tested positive by sarbecoviruses RT-qPCR and at least one of the pan-coronaviruses RT-PCRs) (Supplementary Table 1). The 90 (23 + 67) rectal swab samples positive by sarbecoviruses RT-qPCR included *Rhinolophus* spp. *(R. acuminatus, R. chaseni, R. malayanus, R. microglobosus, R. pusillus*, and *R. shameli*), *Hipposideros* spp. (*H. armiger, H. larvatus*), *Lyroderma lyra, Megaderma spasma*, and *Taphozous melanopogon* as detailed in Extended Data Table 1. We performed Sanger sequencing of the PCR products targeting distinct parts of the RNA-dependent RNA polymerase (RdRp) gene^20,21^ obtained for the 123 (56 + 67) samples that tested positive with at least one of the pan-coronaviruses RT-PCRs.

Phylogenetic analysis of the partial RdRp sequences revealed a diversity alpha- and betacoronaviruses, and seven samples showed signals of potential co-infections (Extended Data Fig. 1): sample RmiSTT521 contained a partial RdRp sequence closely related to alphacoronaviruses (*Rhinacovirus*), and we obtained a near complete genome sequence of a sarbecovirus by high throughput sequencing (see below). The partial RdRp sequences of samples RshSTT288, RshSTT307, RshSTT313, and RshSTT517 clustered with alphacoronaviruses (*Decacovirus*), while samples RshSTT503 and RshSTT523 clustered within betacoronaviruses (*Hibecovirus*). In agreement with the late Ct values obtained in the sarbecovirus RT-qPCR for these samples, we were only able to obtain a partial sarbecovirus RdRp sequence for two of those samples (RshSTT307 and RshSTT313). These three results, supported by sequence data, confirm co-infections involving betacoronaviruses and different alphacoronaviruses. While co-infection is the most likely explanation, we cannot rule out other factors contributing to the signal observed for the four multi-positive samples without sarbecovirus sequence data, such as recombinant viruses. Similarly, two samples among the 56 that tested negative for sarbecoviruses by RT-qPCR but were positive at least one of the pan-coronaviruses RT-PCRs (HlaSTT489 and HlaSTT508) contained two distinct partial RdRp sequences. Phylogenetic analysis showed that one sequence clustered with alphacoronaviruses (*Decacovirus*) and the other with betacoronaviruses (*Hibecovirus*), also indicating likely co-infections. However, the presence of a recombinant virus cannot be excluded.

### Identification of novel bat coronaviruses

We used metatranscriptomic and amplicon-based approaches to generate sequence data from a selection of 50 bat samples that tested positive for sarbecovirus RNA. We obtained 33 full-length and 7 partial genome sequences (Supplementary Table 2 and Supplementary Fig. 1), which were used to infer an initial maximum likelihood phylogenetic tree with publicly available sarbecoviruses sequences (Extended Data Fig. 2). With all the limitations related to recombination in consideration^22^, the phylogenetic analysis revealed the co-circulation of multiple SARS-CoV-2 related viruses in bat populations in Cambodia, most of which were not previously identified or characterized. The seven partial sequences were excluded from further analyses due to their lower quality. Of note, four of them corresponded to non-*Rhinolophus* bat samples (*H. larvatus, L. lyra, M. spasma*, and *T. melanopogon*) with low viral load based on RT-qPCR results. Although extreme care was employed to limit the risk of cross-contamination during sample collection and processing, these results may not reflect productive infections in these bats, and further sampling will be needed to understand whether these viruses can indeed infect these bat species.

One sample (RshSTT564) presented numerous intrahost single nucleotide variants (iSNVs) dispersed throughout the genome at a frequency of ∼0.23 [0.21 to 0.25], suggestive of contamination or co-infection with a closely related virus. This result was reproduced following an independent RNA extraction and sequencing run (Supplementary Fig. 1). Using the intra-host single nucleotide variants (iSNVs) frequency data, we generated an alternate RshSTT564 genome corresponding to the minor variant population (Extended Data Fig. 2). The constellation of mutations in this genome was distinct from all other consensus sequences generated in this work, suggesting that it does not correspond to cross-contamination during library preparation. All virus isolation attempts on both simian (Vero) and bat (Rhileki) cells were unsuccessful.

### Evolutionary history of sarbecoviruses circulating in Cambodia

The evolution of sarbecoviruses is characterized by frequent recombination events^18,23^, making it challenging to reconstruct their evolutionary history using a single phylogeny. We thus used methods implemented in RDP4 to identify recombinant regions within the novel sequences. Based on recombination and phylogenetic analyses (Fig. 1 and Supplementary Fig. 2), we identified distinct genomic patterns among the sarbecoviruses circulating in Cambodia. We categorized these viruses into four groups based on their mosaic genome organization, spike receptor binding domain (RBD) sequence and sampling hosts.

**Fig. 1:**
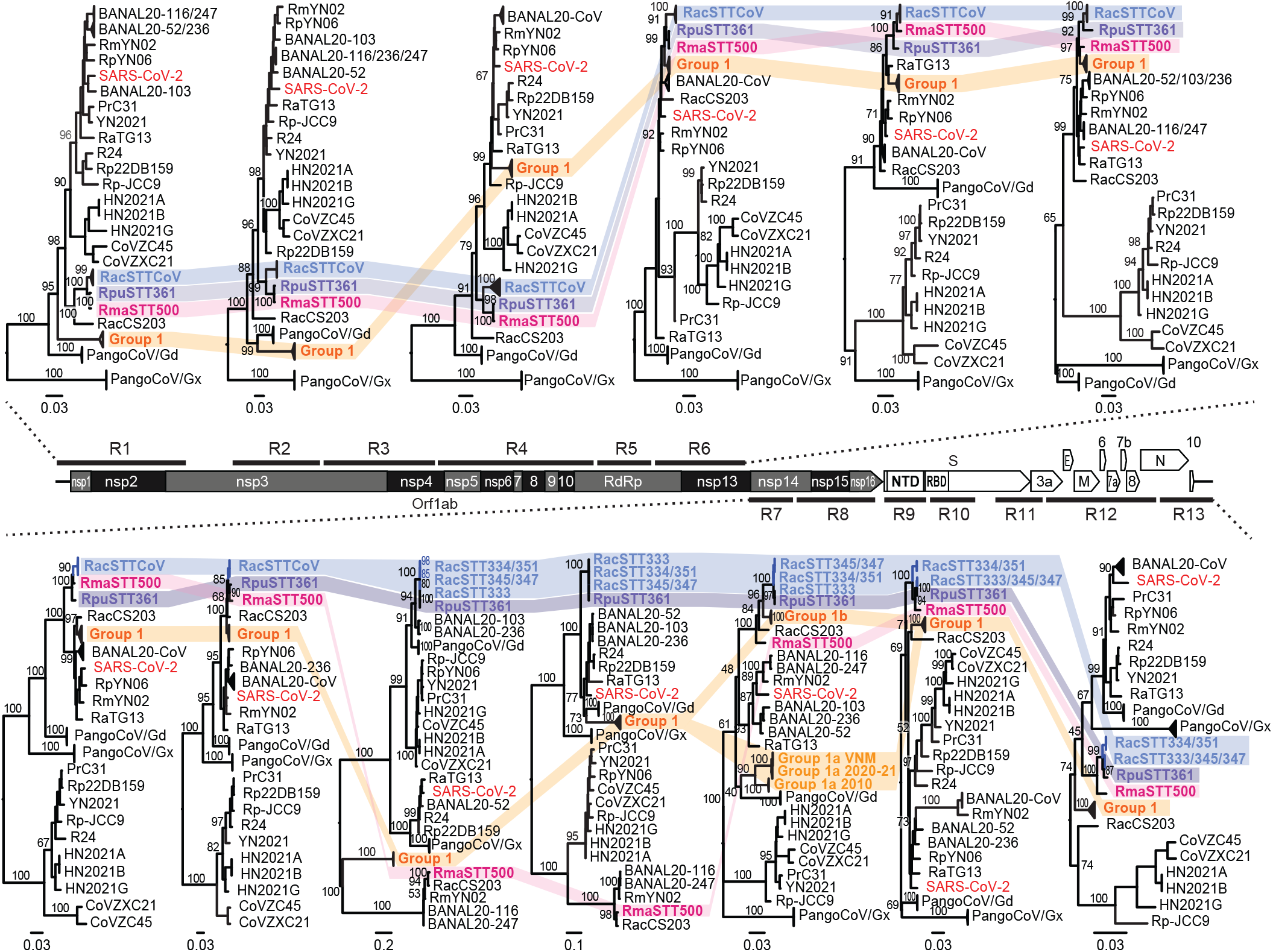
Maximum likelihood phylogenetic trees of the genomic regions identified by recombination analysis. The positions of the 13 genomic regions analyzed are shown on a sarbecovirus genome scheme. Annotated features include ORF1ab with its nonstructural proteins (nsps), the N-terminal domain (NTD) and receptor-binding domain (RBD) of the spike (S) gene, followed by ORF3a, envelope (E), membrane (M), ORF6, ORF7a, ORF7b, ORF8, nucleocapsid (N), and ORF10. The different mosaic patterns are highlighted in colors: group 1 in orange (with lighter and darker shades in region 11 for subgroups 1a and 1b, respectively), group 2 in blue, group 3 in purple and group 4 in pink. Branch support values obtained from 1,000 ultrafast bootstrap replicates are shown on relevant nodes. Branch scale bars represent substitutions per site. Complete (non-collapsed) phylogenies are presented in Supplementary Fig. 2.

The most prevalent sarbecoviruses (designated here as group 1), were found primarily in *R. shameli* samples, the most frequently captured bat species in this study (59% of total bats), as well as in *R. microglobosus* (Extended Data Table 1). These genomes, sampled in 2020, 2021, and 2023, are closely related to two viruses previously detected in samples collected from *R. shameli* bats in the same area in 2010^14^, and to four genomes recently reported from Vietnam in 2022 from *R. affinis* bats^16^. Our analysis identified two mosaic patterns within group 1 (designated as 1a and 1b), with evidence of recombination in the spike S2 domain while the viruses are otherwise monophyletic in all other parts of the genome (Fig. 1, region 11). However, it is unclear whether the subset 1a, which includes the 2010 viruses, are the recombinants, having acquired a portion of S2 from a divergent lineage, or if instead it corresponds to the ancestral sequence of group 1 viruses, with subset 1b having recombined with viruses of the group 2, 3, or 4 lineages. Nevertheless, the absence of a close relative to the divergent S2 sequence suggests that at least one additional sarbecovirus lineage, currently unsampled, circulates or has circulated in recent years in the region. In parallel, group 2, 3, and 4 viruses, which were sampled from *R. acuminatus, R. pusillus*, and *R. malayanus* respectively, constitute a novel subclade within the sarbecovirus phylogeny. These viruses cluster together in most regions of the genome (Fig. 1), with branching variations suggestive of inter-group recombination. Of note, the group 4 virus, together with RaCS203, cluster separately from groups 2 and 3 in the regions of the spike. RaCS203 was identified in Thailand in 2020, and it is also found basal to the viruses sampled in Cambodia in multiple other parts of the genome, further suggesting both common ancestry and genetic exchanges during the evolutionary history of the viruses found in these neighboring countries. Finally, our selective pressure analysis across the full dataset did not identify codon sites under positive selection with different methods.

To better understand the evolutionary histories of the sarbecoviruses co-circulating in Cambodia, we inferred time-measured phylogenies using a Bayesian phylogenetic approach for each of the 13 identified genomic regions as well as for a non-recombining alignment (NRA, see Methods) (Supplementary Fig. 3). In addition, as our data also allow the study of the continuous evolution of group 1 viruses lineage, which spans a 13-year window, we separately studied this subset of viruses, aiming to leverage complete genome data. However, as viruses within this group present evidence of recombination in the spike S2 domain (region 11), we conducted the phylodynamic analysis focusing on the genome truncated from region 11, which corresponds to 95.7% of the complete alignment (Extended Data Fig. 3). We used two datasets: group 1a viruses only, as they appear to have a continuous evolutionary history, and the complete group 1 viruses set that includes 1b viruses with a possibly distinct history. Using these near-complete genome data, the substitution rate estimated for group 1a was 6.82 × 10^−4^ substitutions per site□yr^−1^ [95%HPDs 3.59 ×□10^−4^ – 9.15□×□10^−4^], and slightly faster for the complete group 1 dataset (1.02 × 10^−3^ substitutions per site□yr^−1^, Supplementary Fig. 4). Our estimates are within the range of the mean estimates previously reported for sarbecoviruses using partial (non-recombinant regions) genome data^18^, and comparable to estimates obtained for SARS-CoV-2^24^ or MERS-CoV^18^. These substitution rates may be affected by the small sample size and relatively short time range and may also reflect incomplete purifying selection processes. Similarly, we noted variation in the substitution rates estimates across the 13 non-recombinant regions and the NRA, likely reflecting a combination of factors including alignment length, genomic location and potential functional constraints, or selective pressures (Supplementary Fig. 4).

Despite this variation, our results, both from the phylodynamic and the recombination analysis focusing on divergence times between specific lineages in relevant genomic regions, are compatible with multiple genetic exchanges over time between viral lineages sampled both within and outside Cambodia. Group 3 and group 4 viruses are monophyletic in 5 of the 13 genomic regions and in the NRA, and we estimate a median time to the most recent common ancestor (tMRCA) as recent as 6.5 months prior to the earliest sampling (Fig. 2a, and tMRCA visualized as dates in Supplementary Fig. 3), with short estimates obtained for 4 regions and the NRA. Within the same 5 regions and the NRA, the earliest median tMRCA for group 2 viruses with group 3-4 viruses was estimated at 11.5 years (Fig. 2b), although with greater variation between regions, as expected for a deeper node. Based on regions 9 and 10, corresponding to mostly the spike, we estimated that a common ancestor to the group 4 virus and the RacCS203 virus sampled in Thailand, ∼470km away from the Steung Treng region of Cambodia, cocirculated with these lineages 7.8 to 14 years prior to the sampling [95% highest posterior density HPD 0.5 – 31.2 years] (Fig. 2c). Finally, within group 1, the viruses from Vietnam share a common ancestor with viruses from Cambodia with a median divergence time estimate of July 2020 to January 2021 (using group 1 and 1a respectively) (Extended Data Fig. 3). This suggests that these viruses would have spread across a distance of ∼345 km (between Cambodia’s Steung Treng province and Vietnam’s Quang Tri province) within 1.74 to 2.94 years [95% HPD for group 1] or within 1.64 to 3.27 years [95%HPD for group 1a] (Fig. 2d).

**Fig. 2:**
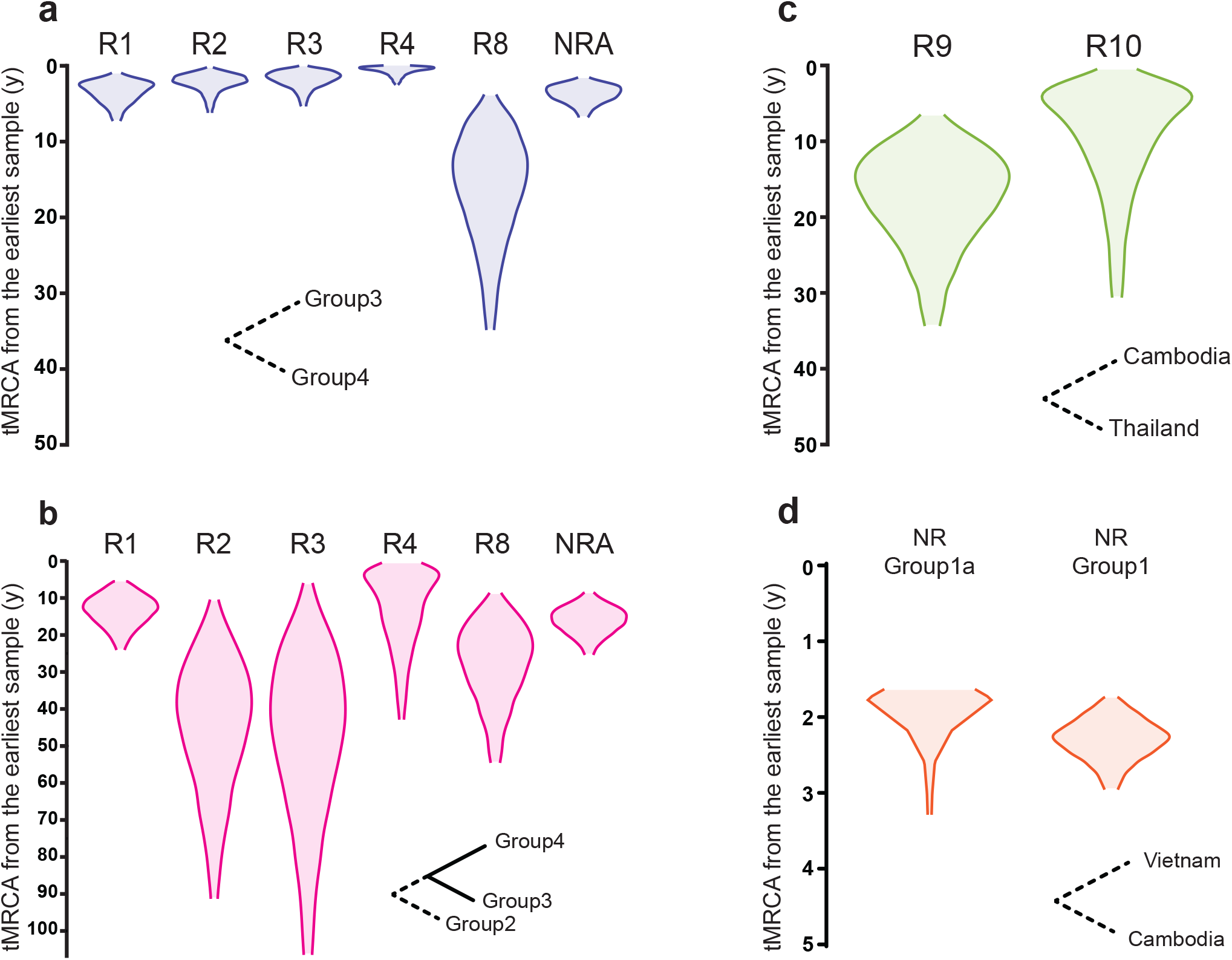
Distribution of the estimated time to the closest inferred ancestor of the viruses from bats sampled in Cambodia, estimated from different regions of the genome. a, Distributions for the time to the closest ancestor to viruses from groups 3 and 4, estimated using regions 1-4, 8 and the non-recombining alignment (NRA). b, Distributions for the time to the closest ancestor to viruses from group 2 and the ancestor to group 3 and 4, estimated using regions 1-4, 8 and the NRA. c, Distributions for the time to the closest ancestor to viruses from group 4 and the RacCS2003 virus sampled in Thailand, estimated from regions 9 and 10. d, Distribution of the time to the closest ancestor to virus from group 1 sampled in Vietnam, and viruses sampled in Cambodia. The violin plots represent the 95% highest posterior density (HPD) intervals of time using the oldest sample of each pair of clades as a reference time point. The corresponding violin plots visualized as calendar dates are presented in each of the Maximum Clade Credibility (MCC) trees in Supplementary Fig. 3.

### Analysis of the spikes and RBDs

We then focused our analysis on the spike protein of the different lineages co-circulating in Cambodia. First, considering that the four groups of viruses identified here were found in distinct bat species, we generated a tanglegram between a spike amino-acid phylogeny (again with the limitations due to recombination in mind) and the angiotensin-converting enzyme 2 (ACE2) amino acid sequences phylogeny of representative bat species, pangolin and human (Extended Data Fig. 4). This comparison revealed a clustering pattern with the 3 bat species associated with group 1 viruses form a distinct clade from the bat species harboring viruses from group 2, 3 and 4.

Next, we used homology modeling and sequence analysis to compare the external subdomain of each novel spike RBD structure to the SARS-CoV-2 RBD structure, and found that they are globally similar with some specific differences (Fig. 3). Group 2-3 RBDs are highly similar to SARS-CoV-2 RBD, with only one amino acid difference among positions identified as contact residues for human ACE2 (hACE2) receptor binding^25^. This type of receptor-binding motif (RBM), also observed in viruses identified in Laos^15^, has been associated with broad ACE2 tropism, including efficient hACE2 binding. In contrast, group 1 viruses RBDs present two to three differences at key contact residues, as well as a shorter loop at the beginning of the RBM (spanning residues S:443-450, SARS-CoV-2 numbering), and a single amino acid deletion in loop 469-491 (Fig. 3). This pattern has been associated with tropism to a narrower range of ACE2 orthologs and found generally incompatible with efficient hACE2 usage. Finally, and consistently with the analysis of the RacCS203 and RmYN02 virus RBDs^8,13^, the group 4 virus displayed marked structural differences with SARS-CoV-2 RBD, including two shorter loops at the N terminal and central regions of the RBM, with the absence of residues involved in a conserved disulfide bond, and variations at most positions identified as contact residues for hACE2 receptor binding (Fig. 3). None of the newly identified viruses harbored a poly-basic (Furin) cleavage site as present at the S1/S2 junction in SARS-CoV-2 (Supplementary Fig. 5). However, the group 4 virus, like viruses from Thailand and China, presents a three amino acid insertion (PVA) at this site.

**Fig. 3:**
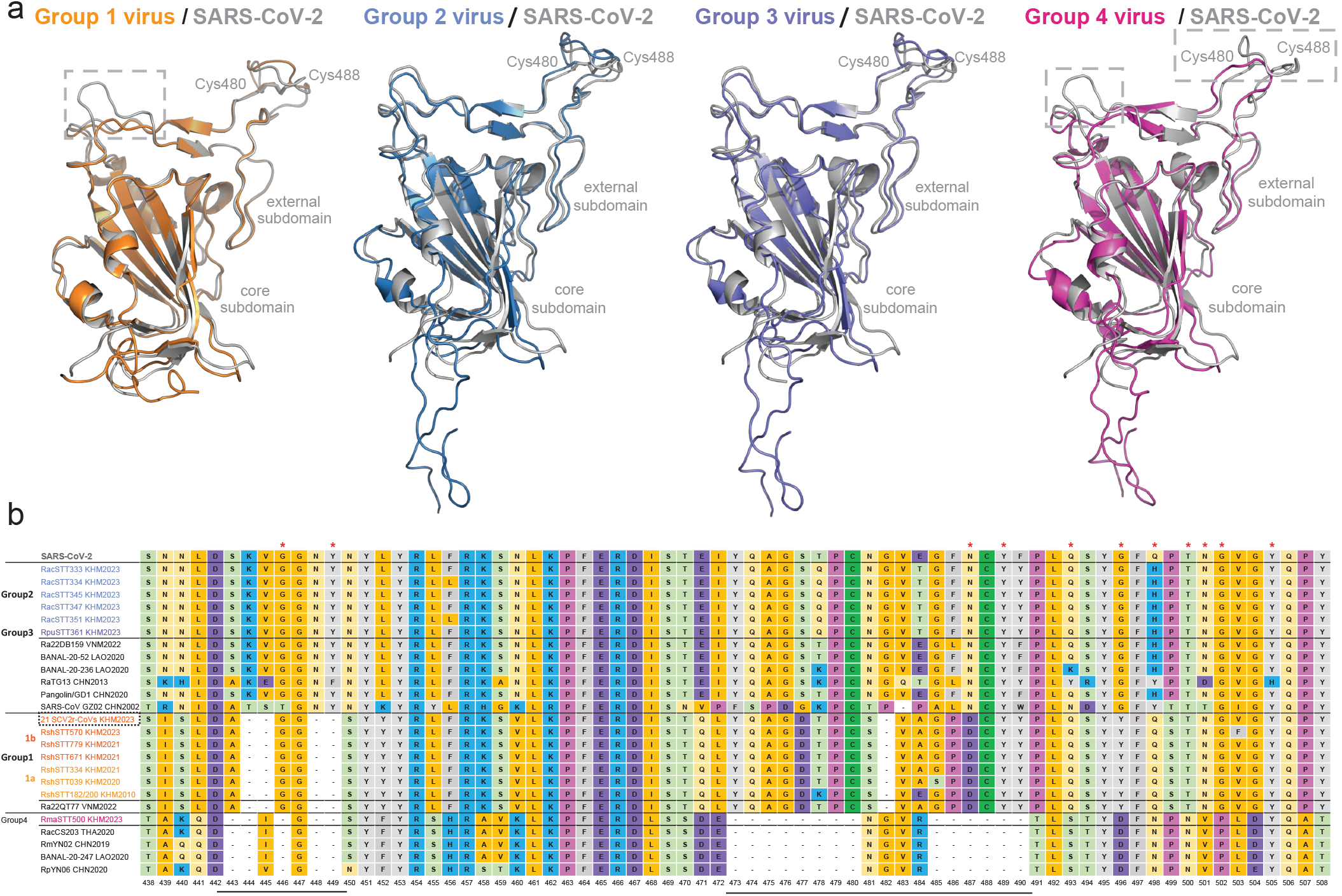
Analysis of the spike receptor-binding domain (RBD) of SARS-CoV-2 related bat viruses sampled in Cambodia. a, Homology modeling of the structure of the RBDs of the 4 groups of viruses from Cambodia compared to SARS-CoV-2 RBD structure (PDB: 6yla.1). The three-dimensional structure of the RBDs of the viruses from Cambodia were modeled using the Swiss-Model program using the most fitting template. The structure of RshSTT182/200 RBD (PDB: 7xbh.1) was used for virus group 1 (orange), the structure of BANAL-20-52 Spike trimer (PDB: 8hxj.1) was used for virus group 2 (light blue) and 3 (dark blue), and the structure coronavirus spike protein (PDB: 8aja.1) was used for virus group 4 (pink). b, Alignment of the receptor binding motif amino acid sequences of selected sarbecoviruses. Residues reported as involved in the interaction between SARS-CoV-2 RBD and the human ACE2 receptor^25^ are marked with a red asterisk (*).

### Functional assays

To further investigate the potential for human infection, we used a lentivirus-based pseudovirus system to assess the ability of representative spikes proteins from group 1 and group 2-3 viruses to mediate entry into cells expressing hACE2, with the SARS-CoV-2 (Wuhan-Hu-1) spike as a control (Fig. 4a). The group 2 virus (RacSTT345) spike was able to mediate entry into cells expressing hACE2 (Fig. 4b), suggesting a lower barrier for spillover. On the other hand, as observed for the 2010 viruses, the recent group 1 spikes (with or without recombination in S2, RshSTT039_2020 (group 1a) and RshSTT671_2021 (group 1b) respectively) were not able to mediate entry into cells expressing hACE2, while they were able to mediate entry in cells expressing *R. shameli* ACE2 (RshACE2) (Fig. 4b,c). Of note, previous work has reported low levels of entry into hACE2-expressing cells using a VSV-based pseudovirus system with the 2010 spike, which could be improved by a single amino-acid mutation^26^. The isolation of a group 1 virus will be crucial to clarify the zoonotic potential of this viral lineage.

**Fig. 4:**
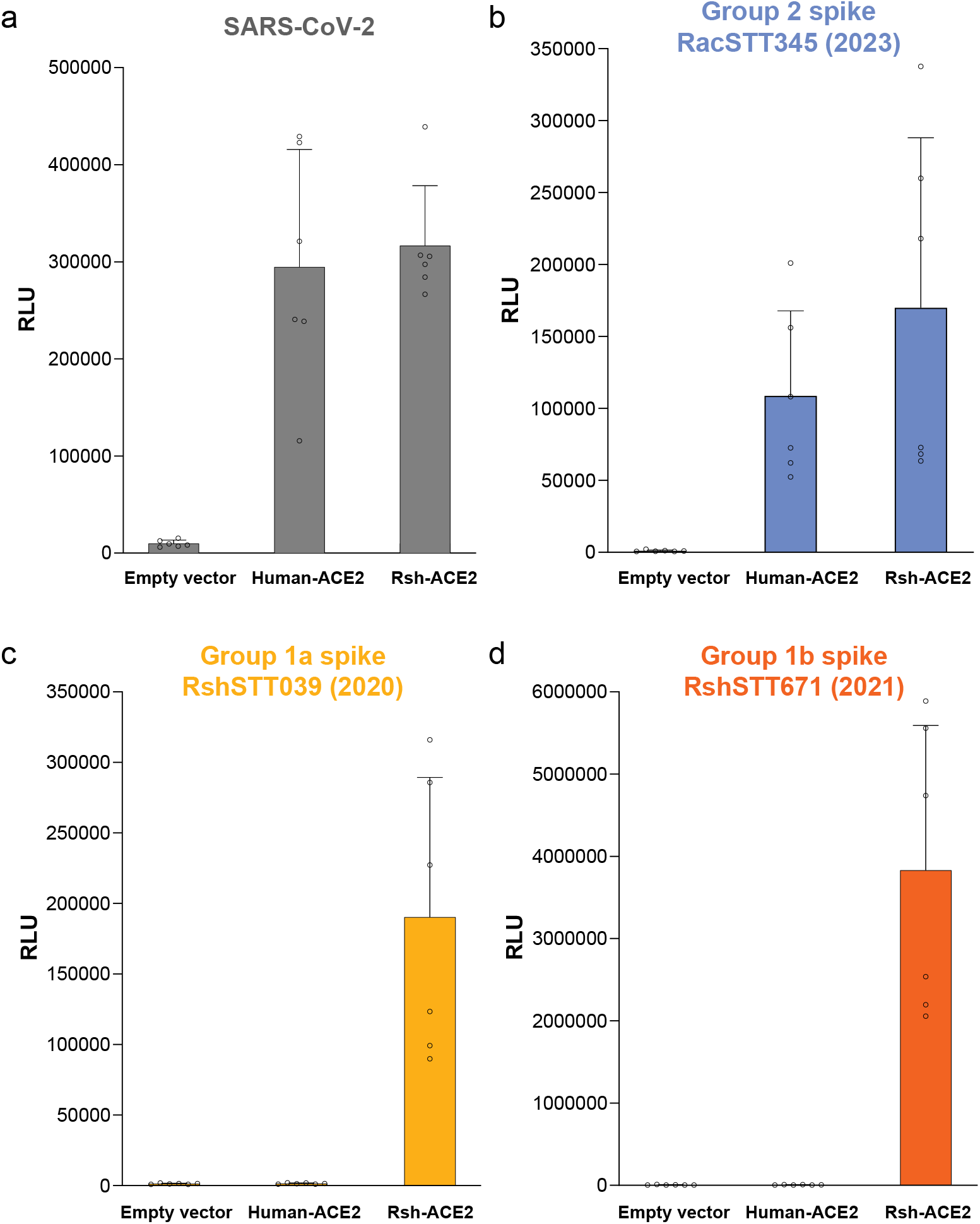
Pseudovirus entry assays. Entry into HEK293T cells transfected with either human ACE2 (hACE2), Rhinolophus shameli ACE2 (RshACE2), or the corresponding empty vector (pLenti-puro) of pseudoviruses expressing the spike of a, SARS-CoV-2, b, group 2 RacSTT345, c, group 1a RshSTT039 and d, group 1b RshSTT671. Data are expressed in relative luminescence units (RLU) measured 48 h post-pseudovirus inoculation and are represented as mean ± standard deviation of technical replicates for two independent experiments (n=6). Source data are provided as a Source Data file.

## Discussion

In this study, we report the longitudinal sampling of bats belonging to nine genera in the Steung Treng region of Cambodia, with a particular focus on *Rhinolophus* species. Using multiple sequencing approaches, we generated 33 novel sarbecovirus genomes, uncovering a much greater viral diversity than previously observed in the country. Through recombination and phylogenetic analyses, we identified distinct genomic patterns and evidence of further, unsampled, diversity. Although these different lineages were found in distinct groups of bat species, all the sarbecoviruses identified in Cambodia appear to share a common evolutionary history, at least in part, as they cluster closely together in certain regions of their genomes in the context of the currently known diversity of sarbecoviruses (Fig. 1). In addition, variations in tree topologies across the different genomic regions are consistent with recombination, - and thus past co-infection of a host - between those lineages circulating in Cambodia.

These genomic exchanges are not confined to lineages found in Cambodia. Viruses recently detected in Vietnam fall within the diversity of group 1 viruses, and share a very recent ancestry, with estimates in the order of two years, despite the geographic distance between the sampling sites. Similarly, a virus identified in Thailand in 2020 (RacCS203) shows close genetic proximity to the viral lineages from Cambodia, branching basal or within their diversity in multiple regions of the genome, evidencing a common ancestry. Furthermore, the group 4 virus exhibits strong evidence of recombination in the spike gene with an ancestor of RacCS203. Together, these findings suggest frequent and relatively long-distance virus migrations have occurred, potentially via bat migrations or transmissions across neighboring populations. This is particularly notable in very recent years for group 1 viruses, possibly facilitated by the dense forest environment that spans Northern Cambodia and Vietnam. Further studies will be necessary to determine whether the ongoing land-use changes in Southeast Asia, which may impact bat habitats, have contributed to these movements, alongside the potential effects of climate change^27^.

Even within more restricted geographic areas, such as the sites surveyed in the Steung Treng province, the habitat-sharing behavior of diverse bat species appears to facilitate viral exchanges, offering opportunities to expand the genetic diversity of viral populations. Interestingly, the four groups of newly detected viruses were each found to be associated with distinct bat species, despite being captured in the same geographical area where they likely share the same or close habitats. While further sampling could alter this observation, the current observation may reflect finer co-roosting habits of these species, or host-virus compatibilities and virus adaptations distinct from spike-ACE2 receptor interactions. Indeed, although ACE2 amino acid phylogeny groups the bat species harboring the sarbecoviruses identified in Cambodia into two broad clusters (Extended Data Fig. 4), the evidence of recombination and functional assay data indicate that intergroup infections can occur. The 2010 group 1 spike has been shown to be able to mediate entry in cells expressing *R. malayanus* ACE2^28^ and we showed here that a group 2 spike is compatible with *R. shameli* ACE2. Isolating these viruses would be key to understanding these potential restrictions. Similarly, further sampling would also help to determine whether the low viral load signal and partial sequence data obtained from several non-*Rhinolophus* bats samples reflect productive infections, as this would come to challenge our understanding of the extent of the host range of these viruses.

Finally, as a crude indicator of emergence risk, we assessed the capacity of spike proteins from group 1 and 2 to mediate entry into cells expressing hACE2. While the recent spikes of group 1 viruses were not compatible with human ACE2 in our pseudovirus system, consistent with previous findings with a spike from 2010^14^, the group 2 spike (RacSTT345) was able to mediate entry, suggesting potential for spillover. Interestingly, this spike was also able to mediate entry into cells expressing RshACE2, and this, with the high similarity with SARS-CoV-2 RBD amino acid sequence, suggests that group 2 and 3 spikes may also be generalists in terms of host tropism^29^. However, there is currently no indication of a significant threat to public health, given the absence of a furin cleavage site. Additionally, immunity induced by SARS-CoV-2 exposure or vaccination may offer cross protection against these viruses, considering the high amino acid similarity of the group 2 and 3 spike RBDs to SARS-CoV-2, reaching up to 96.8% identity—a value comparable to the highest similarity reported for viruses from Laos^15^. Interestingly, the nucleotide-level similarity of group 2 and 3 RBDs is lower, with numerous synonymous mutations compared to SARS-CoV-2, which may reflect host adaptation (at the species level) through codon usage optimization.

This work underscores the extensive genomic diversity of coronaviruses co-circulating in a small area of northern Cambodia and indicates both local and regional lineage migrations over short periods. Expanding surveillance efforts geographically and longitudinally will be crucial to identify high-risk animal-to-human interfaces and track their evolution to strengthen future pandemic preparedness efforts.

### Limitations of the study

Our work has several limitations. First, we were unable to isolate any of the viruses reported here. Successfully isolating or reconstructing these viruses via reverse genetics would allow for a proper assessment of their properties, host range, and a better understanding of their potential emergence risk. Additionally, it would be valuable to evaluate whether sera from individuals vaccinated against COVID-19 or with history of SARS-CoV-2 infection can neutralize these viruses. Second, although it was not the focus of this work, the timeline and frequency of sampling missions did not allow us to accurately capture the true temporal trends and ecological determinants of sarbecoviruses prevalence within these bat populations. Strong variations in the detection frequency over the years may be linked to the seasonality of bat gestation and birth, with synchronized pulses of immunologically naive individuals potentially facilitating the amplification of viral populations^30^. Further sampling efforts will be needed to reliably assess temporal patterns and enhance our understanding of the ecological drivers behind virus fluctuations in these bat communities. Despite this, our study revealed an extensive genetic diversity of sarbecoviruses found in a limited area within the Steung Treng province, with evidence of viral lineages movements. This highlights the need for broader sampling across the country and the region to better capture the extent and dynamics of sarbecoviruses circulation.

## Methods

### Sampling

Eight sampling sessions were conducted on the following dates: August 18-22, 2020; October 12-17, 2020, March 13-17, 2021; August 30-September 3, 2021; November 29-December 7, 2021; March 27-30, 2023; May 2-6, 2023 and June 12-17, 2023. Each mission spanned consecutive nights across four different karst hill locations (Phnom Chab Phleurng, Phnom Chhngauk, Phnom Ka Ngoak, and Phnom Kork Romeas) (Extended Data Fig. 5). Mist nets were set up each evening, either at the entrance of caves or on potential bat flyways and monitored during a consistent 2-hour period per night. Capture, handling, species identification, sampling and release of live bats followed established standard procedures used by the study group and adhered to international standards for animal research^31,32^. For each captured bat, species, sex, age, and reproductive status of bat females were determined on site. Bat species were identified using morphological criteria in the field. The following biological samples were collected from captured bats: blood, oral swab, rectal swab, and wing punch. All field activities were conducted in accordance with appropriate biosafety standards^31^. Personnel wore suitable personal protective equipment, and strict decontamination procedures were applied to all materials and equipment used during sampling and transport. All samples were stored on ice during collection and subsequently in liquid nitrogen in the field, and subsequently transferred to the Institut Pasteur du Cambodge biobank, where they were stored at -80°C. Downstream processing was conducted under biosafety level 3 (BSL-3) conditions.

### RNA extraction and molecular testing

RNAs were extracted from the rectal swab specimens using the Zymo Research Direct-zol RNA MiniPrep kit (Zymo Research, CA, USA). RNA samples were subjected to real-time RT-qPCR^19^ targeting the E and N genes for sarbecoviruses detection, and to two pan-coronaviruses conventional RT-PCRs^20,21^ targeting distinct portions of the RdRp gene.

The amplification products of the positive RT-PCRs were Sanger sequenced (Macrogen, Inc., Seoul, Republic of Korea). The sequences were aligned using MAFFT v7.508^33^ with a subset of selected reference sequences retrieved on GenBank (https://www.ncbi.nlm.nih.gov/) and GISAID (https://www.gisaid.org/). The 294 nt long sequence alignment of the target region of Quan et al^20^ includes 107 sequences from this study and 65 publicly available sequences. The 394 nt long alignment of the target region of Chu et al^21^ (distinct from the Quan et al target) includes 115 sequences from this study and 72 reference sequences.

The species of bats positive for a coronavirus was confirmed via DNA barcoding, using conventional PCR to amplify and Sanger sequence the mitochondrial cytochrome c oxidase subunit 1 (CO1) gene with previously described primers^34,35^, followed by BLAST^36^ analysis. Additionally, the ACE2 of bats of interest were sequenced on an Oxford Nanopore Technology (ONT) GridION using previously described primers^37^.

### Sequencing and genome assembly

We employed a combination of sequencing approaches to attempt to obtain viral genomic sequences from the samples positive for sarbecoviruses by RT-qPCR (Supplementary Fig. 1). For samples collected from *R. shameli* in 2023, a selection was performed based on Cq values <31, geographic location and sampling dates. No-template controls were included at every step from extraction to sequencing to control for contamination, and no contamination was detected.

#### Metatranscriptomic sequencing

Metatranscriptomic sequencing following RNase H-based ribosomal RNA (rRNA) depletion was first utilized on two samples detected in 2020 and early 2021 (RshSTT039 and RshSTT334 respectively) and it was further conducted on two samples detected in 2023 (RshSTT570 and RacSTT351). Briefly, extracted RNA was treated with Turbo DNase (Ambion), before RNase H-based rRNA depletion. After conversion into ds-cDNA, libraries were prepared with the Nextera XT library preparation kit (Illumina) and sequenced on an Illumina NextSeq 500 platform (2×75 cycles).

Metatranscriptomic sequencing was also attempted with other approaches. Samples were subjected to sequence-independent single-primer amplification (SISPA) as previously described^38^, using the SISPA-Primer A (5’-GTTTCCCACTGGAGGATA-(N9)-3’) for cDNA synthesis, and the SISPA-Primer B (5′-GTTTCCCACTGGAGGATA-3′) for amplification and sequenced on an ONT GridION.

Samples were also processed using the SMARTer Stranded Total RNA-Seq kit v3 (Takara Bio, USA) for cDNA synthesis and library preparation, and sequenced on Illumina NovaSeq platform (Macrogen, Inc., Seoul, Republic of Korea).

#### Amplicon based sequencing

Leveraging these novel data and the genomes from 2010 (EPI_ISL_852605), we designed a highly multiplexed PCR amplicon approach using Primal scheme^39^. Sequence gaps of RshSTT039 were subsequently bridged by Sanger sequencing. For samples with viruses divergent from the 2010 virus, additional amplicon sets were designed and used to re-sequence these samples (https://github.com/Simon-LoriereLab/BatCoVCambodia2020-23). Amplicon sequencing was performed on an ONT GridION. RNAs were used to generate cDNA using SuperScript™ III First-Strand Synthesis SuperMix (Invitrogen, Carlsbad, CA, USA) or LunaScript® RT SuperMix Kit (NEB, E3010) and PCR amplicons were generated using the Q5 Hot Start High-Fidelity DNA Polymerase (NEB, M0493S). PCR products were multiplexed using ONT native barcodes and ran in batches of 12 to 24 samples per flow-cell (FLO-MIN106). The resulting reads were base-called and demultiplexed using Guppy (7.0.9) with a minimum quality score of 9.

The spike gene sequences of a selection of viruses identified as group 1 (RshSTT039, RshSTT671, RshSTT610, RshSTT570), and of all viruses of group 2, 3, and 4 were confirmed by Sanger sequencing.

### Bioinformatic analyses

We employed a similar bioinformatic workflow for all metatranscriptomic data analyses^14^. After demultiplexing, reads were subjected to quality trimming and adapters removal; We then performed de novo assembly using the metaspades option of SPAdes v3.14.0 and MEGAHIT v1.2.9 for Illumina data and canu v2.2 for ONT data. Scaffolds were queried against the NCBI non-redundant protein database using DIAMOND v2.0.4. Scaffolds identified as sarbecovirus genomes were then used as template for iterative mapping using CLC Assembly Cell v5.1.0 (Qiagen) for Illumina reads or minimap2 v2.17-r941 for ONT reads. Aligned reads were manually inspected using Integrative Genomics Viewer (IGV)^40^ v2.12.3 (https://software.broadinstitute.org/software/igv/) and consensus sequences were generated using a minimum threshold of 10X read-depth coverage for Illumina and ONT data, except for RshSTT039, where a minimal coverage of 3 reads was used (Supplementary Fig. 1). iSNVs were determined using iVar v1.3.1 for Illumina reads.

The reads from amplicon sequencing were first filtered for minimum length of 100 nt using seqtk v1.3-r106. Reads were then processed using the ARTIC pipeline (https://github.com/artic-network/artic-ncov2019) to map the reads, trim the primers and generate a first consensus sequence. We also used iVar v1.3.1 to generate a second consensus. The consensus sequences were aligned, and potential artifacts were visually inspected. The cleaned consensus sequence was then used as a reference for a second mapping, and a final consensus was generated after primers trimming. All consensus sequences were produced using a minimum of 20X read-depth coverage. iVar v1.3.1 was used for iSNV analysis of a putative co-infection (sample RshSTT564). Coverage statistics of the partial and complete genomes obtained are presented in Supplementary Fig. 1.

### Dataset

Sarbecovirus genomes sequence and metadata available in the GenBank and GISAID EpiCoV databases (epi_set_250326kw, https://epicov.org/epi3/epi_set/250326kw) were retrieved in June 2024. Sequences with large stretches of missing data were removed, and a set of 34 genomes (Supplementary Table 3) were retained and selected for further analysis. We added the 33 complete, one alternate and 7 partial sequences generated here, and sequences were aligned using MAFFT v.7.508^33^. The alignment was manually checked for accuracy using Geneious Prime 2023.0.1, resulting in a final alignment length of 29,991 nt.

### Phylogenetic inferences

All maximum likelihood (ML) phylogenies were inferred using IQ-TREE v2.2.0^41^, with ModelFinder Plus (MFP) to select the best-fit substitution model^42^ and branch support was estimated using the ultrafast bootstrap approximation with 1,000 replicates^43^.

### Recombination analysis

We employed a combination of methods implemented in RDP4 v4.97^44^ (RDP, GENECONV, Chimaera, MaxChi, Bootscan, SisScan and 3SEQ) to detect potential recombination events in the newly generated bat viruses genomes, and conservatively considered recombination signals detected by at least four methods. The coordinates of the detected breakpoints were used to partition the alignment, resulting in 13 regions. Of note, within these partitions, some sarbecoviruses not sampled in Cambodia also presented evidence of recombination (shown in grey in Supplementary Fig. 2) but were kept in these alignments as they were not the focus of the analyses.

Following the approach described in Boni et al^18^, we also used the results of the recombination analysis (here applied to all the genomes) to generate a NRA by keeping only the major non-recombinant region of each genome, masking minor recombination regions. Four sequences (PrC31, Rp22DB159, YN2021, and HN2021B) were excluded as they displayed recombination signals across their entire genomes. The novel alignment was reanalyzed with RDP4, resulting in an NRA containing 63 genomes.

Finally, we further analyzed the recombination signal detected in region R11 (spike C-terminal domain) within the highly related group 1 genomes, as no close relative corresponding to the donor parent lineage could be identified. To verify that the recombination signals detected could not be due to other processes such as strong selective pressures, we analyzed the nature and distribution of substitutions across the genomes of all group 1 viruses by comparing the 26 genome sequences collected in Cambodia and four from Vietnam to the two earliest sequences detected in 2010. In agreement with the recombination signal, the C-terminal region of the spike appeared enriched in changes compared to the rest of the genome, but mostly with synonymous mutations, a profile more compatible with the recombination hypothesis.

### Bayesian divergence time and evolutionary rate estimation

We used the 13 mosaic regions, the NRA, and non-recombinant regions of group 1 and group 1a for phylodynamic inferences. To examine temporal signal in each dataset, we first performed a root-to-tip regression analysis using TempEst v.1.5.3^45^ and the corresponding ML phylogenies. While all datasets showed temporal signal when selecting the root position that maximized *R*^2^, the signal was only strong for the group 1 viruses phylogenies, with and without the inclusion of the genomes sampled in 2010. We thus also used a marginal likelihood estimation procedure called BETS^46^ (Bayesian Evaluation of Temporal Signal) to further assess the temporal structure. We compared the statistical fit of two molecular clock models, strict and uncorrelated relaxed clock with an underlying lognormal distribution, and two configurations of sampling times: the correct sampling times and no sampling times. The BETS analyses provided positive (log) Bayes factors for the models with real sampling times over those with no sampling times, allowing us to confirm that all our datasets present temporal signal (Supplementary Table 4). In addition, the marginal likelihood and positive Bayes factor results from the BETS analysis indicated a better performance of the relaxed over the strict clock model.

We performed Bayesian phylodynamic analysis on each dataset using BEAST v.1.10.4^47^, with the GTR + I + G substitution model, and an uncorrelated lognormal relaxed clock, testing different model combinations of coalescent tree priors. We used posterior analyses with path (PS) and stepping-stone (SS) sampling methods to identify the best-fit demographic model (constant size, exponential growth, Bayesian Skygrid, Skyline and skyride priors). MCMC samplings were run sufficiently long to ensure effective sampling sizes (ESS) > 200.

According to the PS/SS analysis, the Bayesian Skyline plot was the best-fitting model for all datasets (Supplementary Table 5), except for the R11 dataset, which was better fitted by the Bayesian Skygrid model, and for group 1a dataset, better fitted by the Bayesian Skygrid model.

We used Tracer v.1.7.1 to summarize the evolutionary parameters, including the estimates of the substitution rate and the tMRCA for nodes of interest (Fig. 2, Supplementary Fig. 4) and compared them to published estimates of coronaviruses. We reran a selection of BEAST analyses after excluding sequences identified as recombinant and did not observe differences in parameters estimates. The final trees were summarized using Tree Annotator v.1.10.4 and visualized using FigTree v.1.4.7 (Supplementary Fig. 2).

### Selection analysis

The non-recombinant coding sequences (after removal of region 11, where recombination was identified) of the 32 sarbecoviruses genomes assigned to group 1 were used to perform selection analysis with DATAMONKEY (www.datamonkey.org), and the fixed effects likelihood (FEL), single-likelihood ancestor counting (SLAC), and fast unconstrained Bayesian approximation (FUBAR) methods. Significance level was set with p-value threshold of 0.1 for FEL and SLAC and with posterior probability of 0.9 for FUBAR. Site-by-site estimation of selection pressure was performed using the “dN-dS” test statistic to assess selection types on individual codons. A site detected by more than two algorithms was considered under selection.

Selective pressures were also measured with the non-synonymous to synonymous substitution rate ratio ω (dN/dS), where ω = 1, ω < 1 and ω > 1 were assumed to correspond to neutral, purifying, and positive selection, respectively. The ω ratio was estimated by the maximum-likelihood method implemented in EasyCODEML^48^, using four pairs of models (M0-M3, M1a-M2a, M7-M8 and M8-M8a).

### Structure modeling

The three-dimensional structures of the spike RBD of viruses identified in Cambodia was modeled using the SWISS-MODEL program^49^ and compared to the structure of the SARS-CoV-2 RBD (PDB: 6yla.1). The structure of RshSTT182/200 RBD (PDB: 7xbh.1) was used for virus group 1 (RshSTT039: 1a and RshSTT671: 1b), BANAL-20-52 Spike trimer (PDB: 8hxj.1) for virus group 2 (RacSTT345) and 3 (RpuSTT361), and the SARS-CoV spike protein (PDB: 8aja.1) for virus group 4 (RmaSTT500).

### Cells and virus isolation

HEK293T (Sigma) and Vero (ATCC CCL-81) cells were maintained in Dulbecco’s modified Eagle medium (DMEM, Sigma-Aldrich) supplemented with 10% fetal bovine serum (FCS, Gibco) and 1% penicillin-streptomycin (Pen-Strep, Gibco). Rhileki (*R. lepidus*) cells were maintained in DMEM supplemented with 10% FCS, 1% nonessential amino acids (Gibco), 1 mM sodium pyruvate (Gibco), and 100 U/mL Pen-Strep. All cell lines were grown at 37°C and 5% CO_2_.

Virus isolation was conducted in BSL-3 by inoculating in parallel Vero and Rhileki cells in parallel, followed by daily observation for cytopathic effect for 6 days. At the end of the observation period, cell culture supernatants were harvested and centrifuged to remove cellular debris. To screen for the presence of sarbecoviruses, a pan-sarbecovirus RT-qPCR^19^ assay was applied.

### Pseudovirus entry assay

Lentivirus pseudoparticles displaying different spike proteins of interest were produced using the system described by the Bloom laboratory^50^. The following reagent was obtained through BEI Resources, NIAID, NIH: SARS-Related Coronavirus 2, Wuhan-Hu-1 Spike-Pseudotyped Lentiviral Kit, NR-52948, kindly contributed by Alejandro B. Balazs and Jesse D. Bloom.

The sequences corresponding to the selected spikes deleted of the last 21 amino-acids and codon-optimized for expression in human cells were de novo synthesized (GeneArt, Life Technologies), and cloned into the pHDM expression plasmid from the lentiviral kit. The sequence of each construct was verified by Sanger sequencing.

Briefly, HEK293T cells were seeded in 10 cm dishes. The following day, the cells were co-transfected with 10 μg of pHAGE-CMV-Luc2-IRES-ZsGreen-W (NR-52516), 3.33 μg each of the helper plasmids HDM-Hgpm2 (NR-52717), HDMtat1b (NR-52518), and pRC-CMV-Rev1b (NR-52519), and 5 μg of a spike-expressing plasmid encoding either the SARS-CoV-2 spike or the selected spikes from virus group 1 (RshSTT200, RshSTT039, or RshSTT671) or from group 2 (RacSTT345). The transfection was performed using Lipofectamine 3000 (Invitrogen) according to the manufacturer’s protocol. The supernatants were harvested 72 hours after transfection, clarified by centrifugation, aliquoted, and then stored at -80 °C.

To assay entry, HEK293T were seeded in 96-wells plates one day prior to transfection with either pLenti-puro-RshACE2, pHAGE2-EF1aInt-ACE2-WT (NR52512) or pLenti-puro (empty) using Lipofectamine 3000 (Invitrogen) according to the manufacturer’s protocol. The day after transfection, media was removed and cells were transduced with pseudoparticles expressing one of the spikes with 5□µg/ml of polybrene transfection reagent (Merck-Millipore) in a final volume of 150□µl. Three days later, an equal volume of Bright Glo reagent (Promega) was added and mixed by pipetting. After 10□min of incubation, quantification was done with a Centro XS LB 960 luminometer (Berthold technologies). Two independent replicates were performed.

## Data availability

The raw sequencing data and the partial RdRp sequences were deposited on the European Nucleotide Archive (ENA) under BioProject PRJEB86850^51^. Assembled genomes were deposited on the GISAID EpiCoV database with ID EPI_SET_250414kp (https://doi.org/10.55876/gis8.250414kp) and the COI and ACE2 sequences deposited on ENA under BioProject PRJEB87055. Primers designed in this study are available at https://github.com/Simon-LoriereLab/BatCoVCambodia2020-23.

## Code availability

## Acknowledgements

The investigators thank everyone involved in zoonotic virus surveillance and investigation in the Kingdom of Cambodia, especially the Government of Cambodia for granting permission to conduct this work. We thank the teams and direction from the Department of Wildlife and Biodiversity, Forestry Administration and General Directorate of Animal Health and Production, Ministry of Agriculture, Forestry and Fisheries and Department of Biodiversity, Ministry of Environment. We greatly thank the field and diagnostic teams of the Virology Unit, Institut Pasteur du Cambodge for all their hard work and dedication, and Dr Hélène Guis for her contribution to the supervision of collaborative projects coordinated by CIRAD at Institut Pasteur du Cambodge and her scientific input in veterinary medicine.

We gratefully acknowledge all data contributors, i.e., the Authors and their Originating laboratories responsible for obtaining the specimens, and their Submitting laboratories for generating the genetic sequence and metadata and sharing via the GISAID Initiative, upon which part of this research is based.

This study was supported by the BCOMING project (Horizon Europe project 101059483), funded by the European Union, the RhinoKHoV project funded by MUSE (MUSE-16297), the ZooCoV project funded by the French Research Agency (ANR-20-COVI-000), the Région Occitanie and Pasteur Foundation Asia. The E.S.-L. laboratory is funded by Institut Pasteur, the INCEPTION program (Investissements d’Avenir grant ANR-16-CONV-0005), the Ixcore foundation for research, the French Government’s Labex IBEID (ANR-10-LABX-62-IBEID), the HERA project DURABLE (grant no 101102733) and the NIH PICREID (grant no U01AI151758). This work used the computational and storage services (Maestro cluster) provided by the IT department at Institut Pasteur, Paris.

## Author information

V.C. and J.C. coordinated the projects.

S.L., S.T., D.C. and V.C. coordinated with central and local authorities.

V.D., J.C. and E.S-L. designed and supervised research.

G.JD.S., P.D. and E.A.K. provided key material or scientific input.

J.G., T.H., V.C. and V.H. collected samples.

TP.O. and L.P. processed the samples.

H.A. conducted virus isolation.

M.P. and E.S-L. performed initial sequencing and genome assembly

TP.O. performed genome assembly and annotation.

TP.O., A.B. and E.S-L. performed genome analysis and interpretation.

TP.O. performed homology modeling.

D.D., R.R.G.M., TP.O. and M.P. performed the functional assays.

TP.O. and E.S-L. wrote the paper with contributions from all authors.

### Ethics declarations

The study followed all relevant institutional and national guidelines for animal care and use. A study agreement was obtained from the Forestry Administration who participated in the field investigations and oversaw all aspects of the study for active sampling, as no animal ethics committee exists in Cambodia. All bat captures, handling and sampling were conducted with the participation of officers of the Forestry Administration of Cambodia under a MoU signed between Institut Pasteur du Cambodge and the Ministry of Agriculture, Forestry and Fisheries. Animal capture and handling were in accordance with the guidelines approved by the American Society of Mammalogists^31^, in addition to the requirements of the statutory study permission provided by the national authority responsible for wildlife research, i.e.: the Forestry Administration of the Cambodian Ministry of Agriculture, Forest and Fisheries. The experimental protocol was approved by the Ethical Committee of VetAgro-Sup in Lyon, France (Comité d’Ethique de VetAgroSup n°18, Avis 2301).

## Additional information Extended data

**Extended Data Fig. 1**: Maximum likelihood phylogenetic trees of partial RdRp gene sequences using novel and publicly available data. a, Phylogenetic tree inferred using partial RdRp gene sequences (n= 172, alignment length 294 bp) corresponding to the target of the Quan et al RT-PCR^20^. b, Phylogenetic tree inferred using partial RdRp gene sequences (n=187, alignment length 394 bp) corresponding to the target of the Chu et al RT-PCR^21^. Sequences from viruses identified in Cambodia are noted with a triangle for single signal of infection, and with a red circle for sample with signal of multiple infections. Branch support values□at relevant nodes are shown. Scale bar indicates nucleotide substitutions per site. The classification of coronaviruses was based on the International Committee on Taxonomy of Viruses (https://ictv.global/report/chapter/coronaviridae/coronaviridae).

**Extended Data Fig. 2**: Phylogenetic analysis of SARS-CoV-2 related viruses. The maximum likelihood phylogeny was estimated from complete betacoronavirus genome sequences (n=75) using IQ-TREE with 1000 ultrafast bootstrap replicates. The sarbecovirus lineages genomes identified in Cambodia are colored according to groups identified based on their mosaic genome organization, spike receptor binding domain (RBD) sequence and sampling hosts. Scale bar indicates nucleotide substitutions per site. Of note, recombination has been shown to be prevalent in the evolutionary history of sarbecoviruses, and the topology of this tree should therefore be interpreted with caution. *R*. for *Rhinolophus, H*. for *Hipposideros, L*. for *Lyroderma, T*. for *Taphozous*, and *M*. for *Megaderma*. Partial viral genome sequences are denoted with an asterisk (*).

**Extended Data Fig. 3**: Maximum Clade Credibility trees inferred for the non-recombinant regions of a, group 1a and b, group 1a. Node labels are posterior probabilities indicating support for the node. The violin plot indicates the 95% highest posterior density (HPD) interval distribution for the estimated time to the most recent common ancestor (tMRCA) for the clade of viruses sampled in Vietnam and the closest ancestor sampled in Cambodia. Group 1a viruses are shown in light orange, and group 1b viruses in dark orange.

**Extended Data Fig. 4**: Spike proteins and ACE2 amino acid sequences phylogenies. The left maximum likelihood phylogeny was inferred using the amino acid sequences of the sarbecoviruses spikes (n = 67). Of note, recombination has been shown to be prevalent in the history of sarbecoviruses, and the topology of this tree should be interpreted with caution. The sarbecovirus lineages genomes identified in Cambodia are color-coded according to groups identified based on their mosaic genome organization, spike receptor binding domain (RBD) sequence and sampling hosts: group 1 in orange, group 2 in blue, group 3 in purple and group 4 in pink. The maximum likelihood phylogeny on the right was inferred using the ACE2 amino acid sequences (n=83) from different bat species, pangolin and human. The ACE2 sequences from which sarbecovirus sequence data was obtained are colored using the same code. The scale bars indicate amino-acid substitutions per site. The shaded boxes link the spike sequences groups to the ACE2 sequence of the bat species from which the viruses were sampled in this study.

**Extended Data Fig. 5**: Map of the four karst hills (Phnom Chab Phleurng, Phnom Chhngauk, Phnom Ka Ngoak, and Phnom Kork Romeas) in Steung Treng, Cambodia where sampling missions were conducted.

## Extended Data Table Legends

Extended Data Table 1: Bat species captured by date of collection and tested by sarbecovirus real-time RT-qPCR^19^ and pan-coronavirus RT-PCRs^20,21^, at the four sites during active, longitudinal surveillance in Steung Treng, Cambodia, between 2020 and 2023.

## Supplementary information

### Supplementary Data Figure Legends

**Supplementary Fig. 1**: Genome coverage of 42 bat SARS-CoV-2-related viruses detected in Cambodia. a, Coverage values are the combination of the sequence data obtained with the different approaches. b, Intrahost single nucleotide variant (iSNV) percentages and sequencing coverage depth are plotted relative to genome position, with a minimum read depth threshold indicated as a dashed line.

Group 1 samples were sequenced using the RshCoV amplicon primer set (AmpliSeq). Groups 2-4 were sequenced using a combination of RshCoV, RacCoV, RpuCoV, and RmaCoV AmpliSeqs, along with sequence-independent single-primer amplification (SISPA) and metatranscriptomic methods. Note: For sample RshSTT039, a coverage depth of at least 3 was used at nucleotide positions 7147-7351 and 9856-9867. The red line shows coverage generated by combining BAM files from untargeted and targeted sequencing. Data represent: Group 1 (Nextera-XT + RshCoV), Group 2 (SISPA + RshCoV + RacCoV), Group 3 (SISPA + RshCoV + RpuCoV), and Group 4 (SISPA + RshCoV + RmaCoV). Nextera-XT: Untargeted sequencing with Nextera-XT kit, SISPA: Untargeted SISPA sequencing, RshCoV, RacCoV, RpuCoV, RmaCoV: Primer sets used for amplicon sequencing for groups 1, 2, 3 and 4, respectively.

**Supplementary Fig. 2**: Maximum likelihood phylogenetic trees of each of the genomic regions identified by recombination analysis. The positions of the 13 genomic regions analyzed are shown on a sarbecovirus genome scheme. Annotated features include ORF1ab with its nonstructural proteins (nsps), the N-terminal domain (NTD) and receptor-binding domain (RBD) of the spike (S) gene, followed by ORF3a, envelope (E), membrane (M), ORF6, ORF7a, ORF7b, ORF8, nucleocapsid (N), and ORF10. The different mosaic patterns are highlighted in colors: group 1 in orange (with lighter and darker shades in region 11 for subgroups 1a and 1b, respectively), group 2 in blue, group 3 in purple and group 4 in pink. Branch support values obtained from 1,000 ultrafast bootstrap replicates are shown on relevant nodes. Branch scale bars represent substitutions per site and the trees are midpoint rooted for clarity.

**Supplementary Fig. 3**: Maximum clade credibility (MCC) trees inferred for each of the 13 genomic regions and the NRA. Posterior probabilities are indicated for relevant nodes. Violin plots denote probability distributions for the estimated time to the most recent common ancestor (tMRCA) of clades of interest. The tips names of the viruses sampled in Cambodia are colored according to the group identified based on their mosaic genome organization, spike receptor binding domain (RBD) sequence and sampling hosts: group 1 in orange (with lighter and darker shades in region 11 for subgroups 1a and 1b respectively), group 2 in blue, group 3 in purple and group 4 in pink.

**Supplementary Fig. 4**: Substitution rate estimates across genomic regions and virus groups. The mean and 95% highest posterior density (HPD) of the substitution rate estimates obtained with the best model tested for the 13 regions, the NRA, and the near complete genomes for group 1 and subgroup 1a are presented in black with a diamond symbol. For comparison, mean substitution rates and 95% HPD estimated in previous studies are presented in grey with a circle symbol. The SARS-CoV, hCoV-OC43, MERS-CoV, and NRR1, 2 and 3 estimates are from Boni et al.^18^. The SARS-CoV-2 estimate is from Markov et al.^24^.

**Supplementary Fig. 5**: Comparison of the spike protein furin-like S1/S2 cleavage site sequences in the spike proteins of SARS-CoV-2, SARS-CoV-2-related viruses detected in Cambodia, and other selected representative sarbecoviruses. The alignment highlights the presence or absence of a furin-like cleavage site, as well as sequence variation in this region. Sequences are grouped by phylogenetic relatedness and colored according to virus origin or host species where applicable.

## Supplementary Table Legends

**Supplementary Table S1:** Summary of 146 positive and 29 negative bat samples tested by Sarbecovirus RT-qPCR^19^ and Pan-coronaviruses RT-PCRs^20,21^, used for species identification and ACE2 sequencing.

**Supplementary Table S2:** List of the 50 SARS-CoV-2 related virus positive bats samples processed for sequencing. NA: Not available. RdRp: RNA-dependent RNA polymerase. The sarbecovirus RT-qPCR corresponds to the E and N genes targets from Corman et al^19^.

**Supplementary Table S3:** List of publicly available sequences used in this study.

**Supplementary Table S4:** Evaluation of temporal signal and clock models comparisons. We tested two molecular clock models. SC: Strict clock, UCLN: uncorrelated relaxed clock with lognormal distribution. Log-marginal likelihood estimates using the generalized stepping stone sampling (GSS) model selection approach are presented for dataset using the sample dates, and the same dataset analyzed without dates, resulting in positive Bayes factors and indicating that there was temporal signal in the data.

**Supplementary Table S5**: Coalescent tree priors comparisons. The log marginal likelihood values were estimated using both path sampling (PS) and stepping-stone sampling (SS), and the values highlighted in bold indicate the best supported model from log Bayes factor comparisons.

