## Supplementary figures and images for "Characterization and evolutionary history of novel SARS-CoV-2-related viruses in bats from Cambodia"

### Extended_Data_Figures

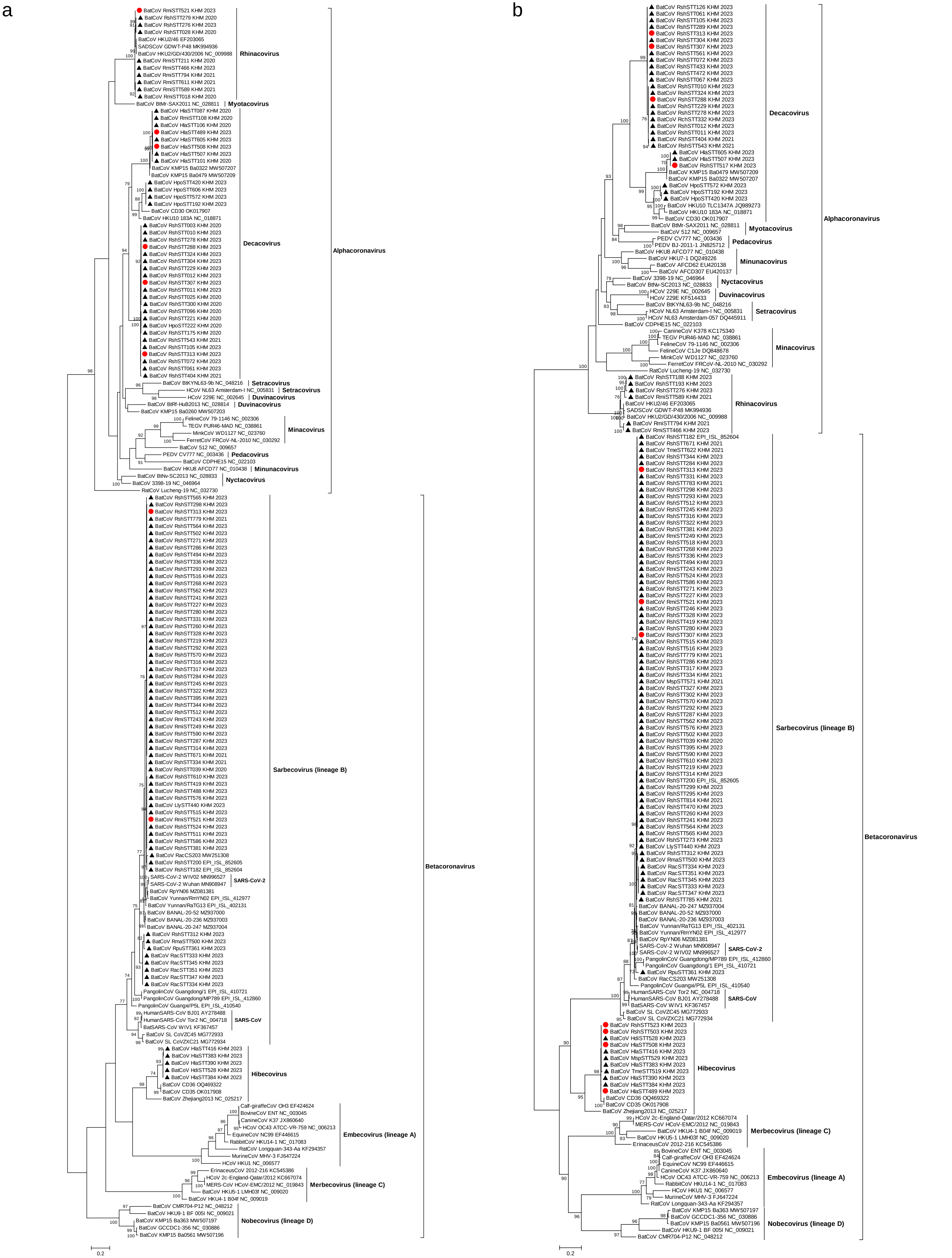

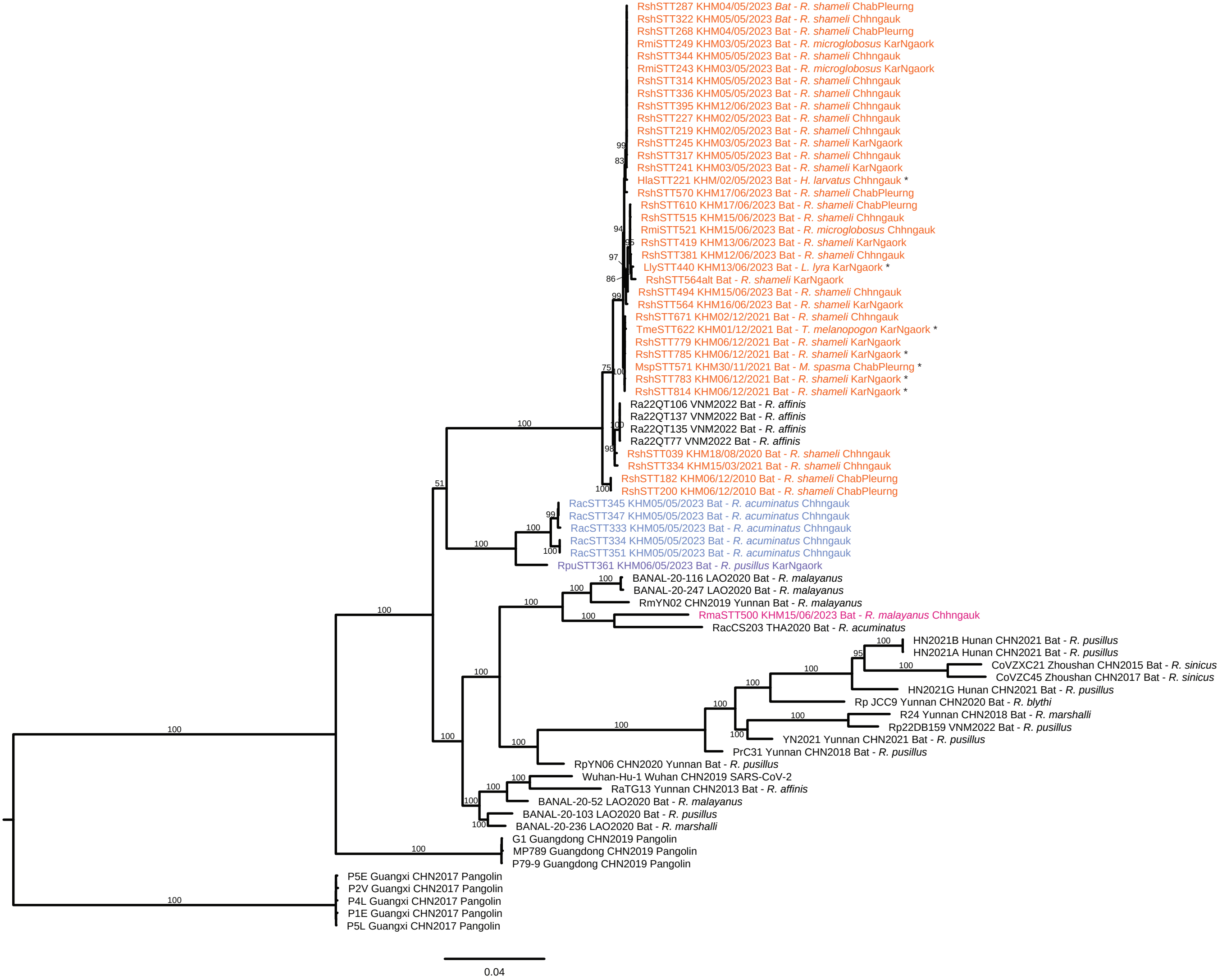

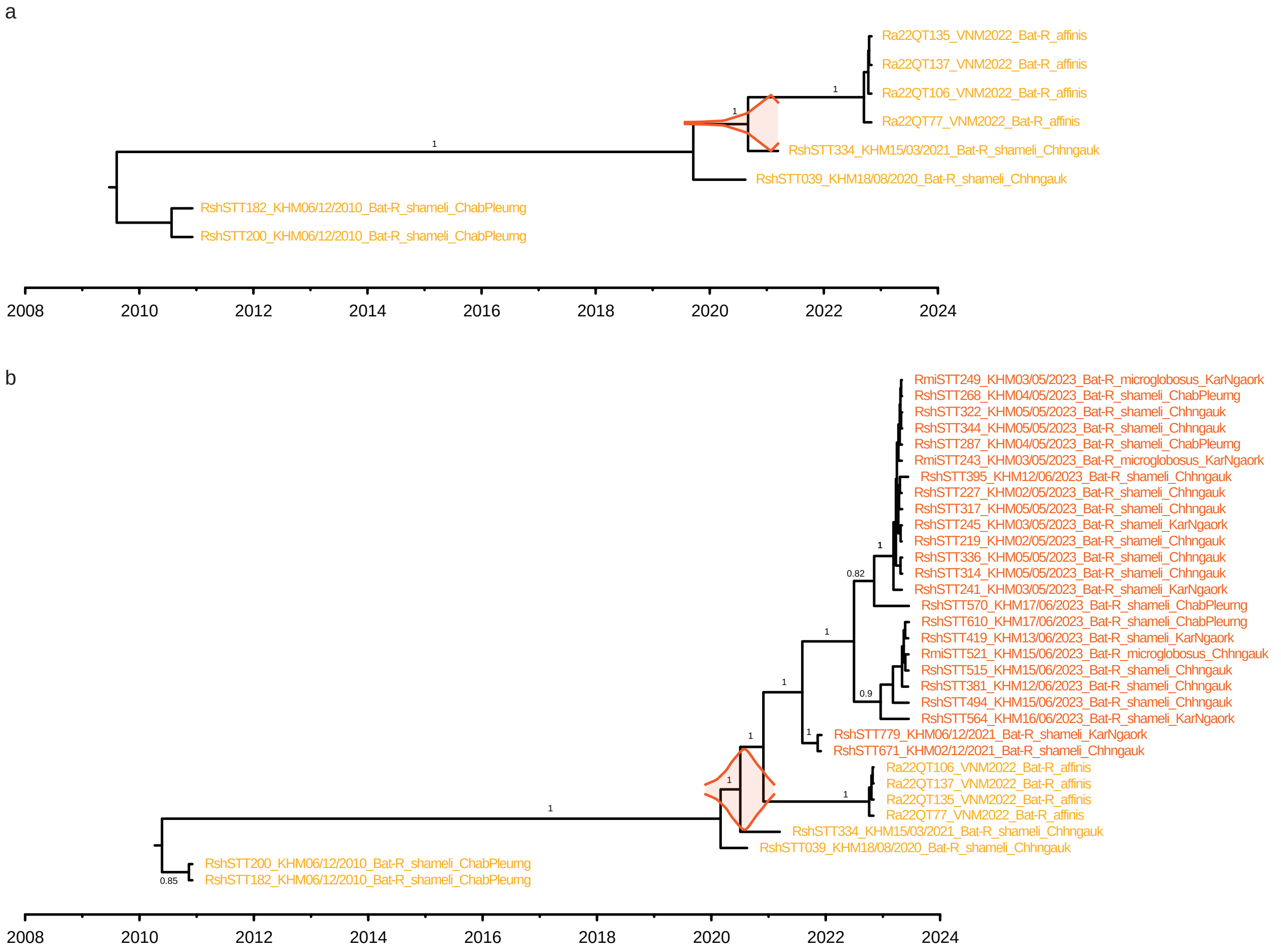

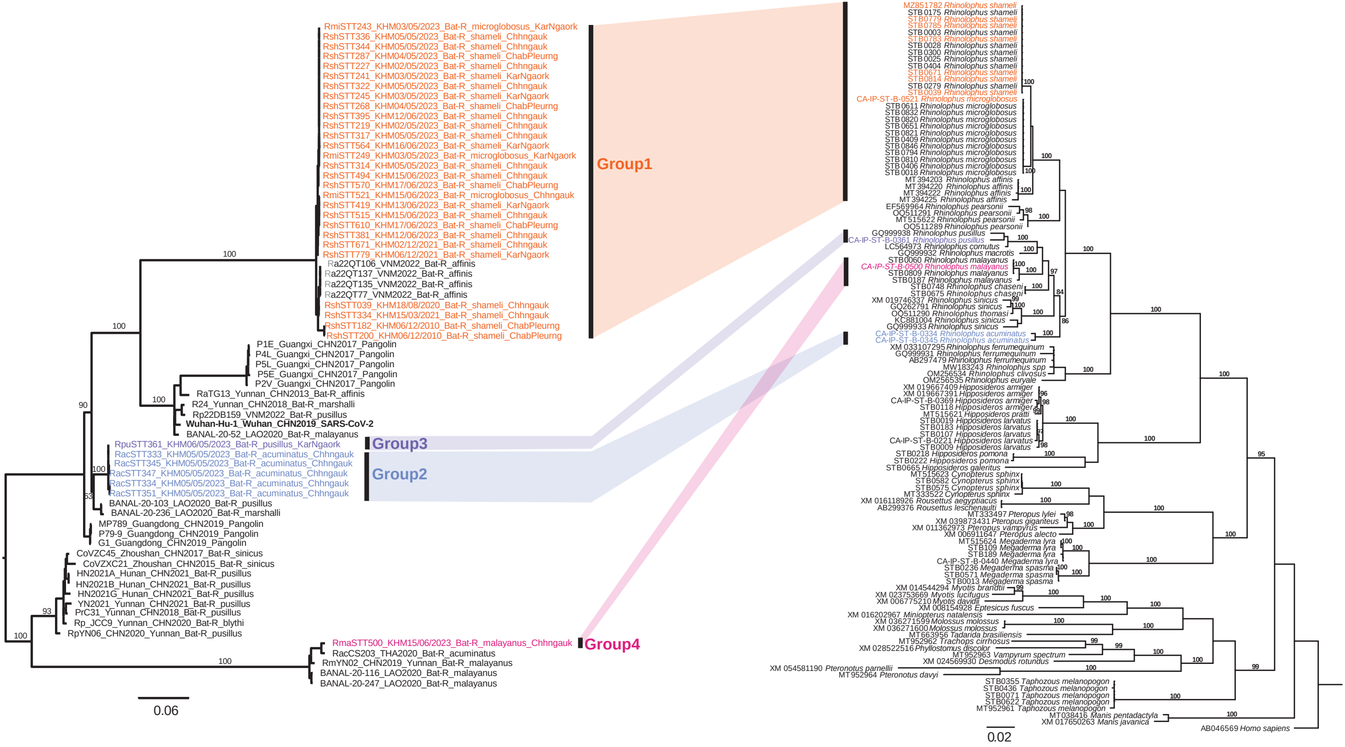

Extended data fig. 5

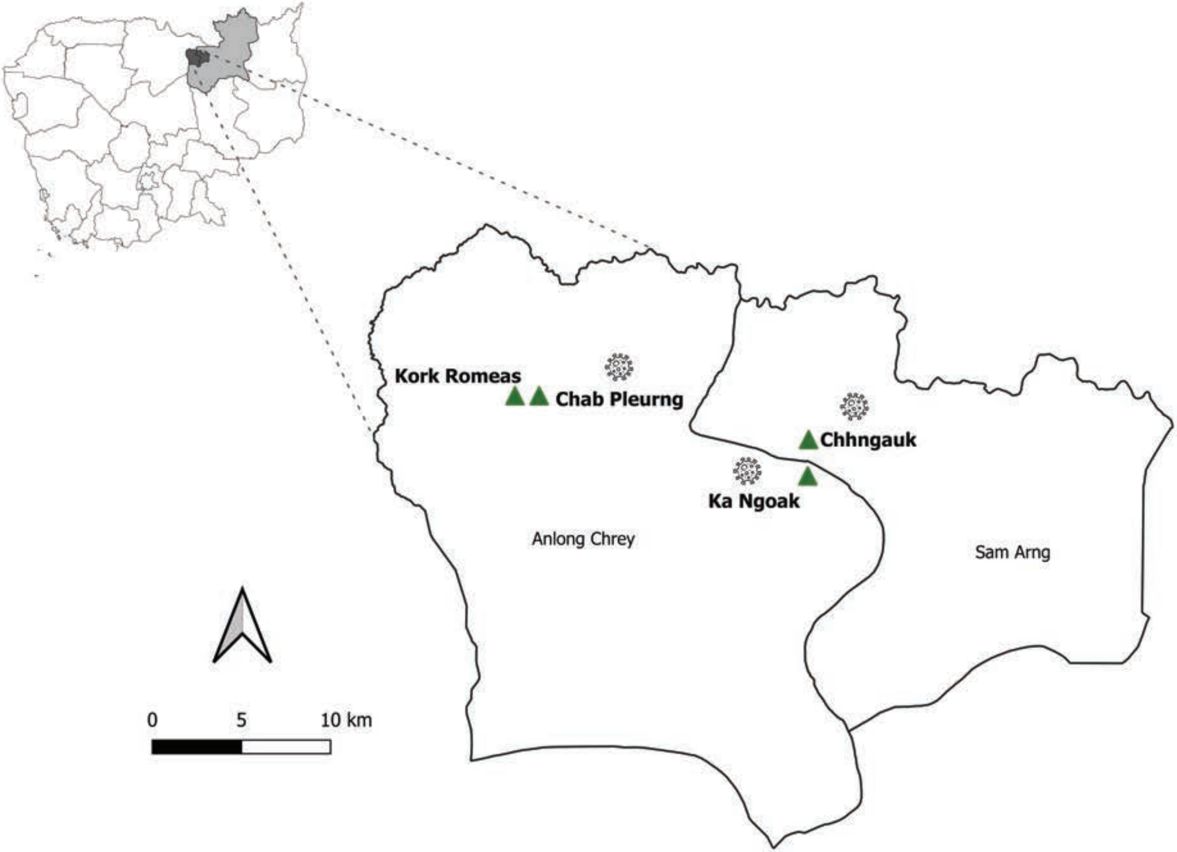

### Supplementary_Figures

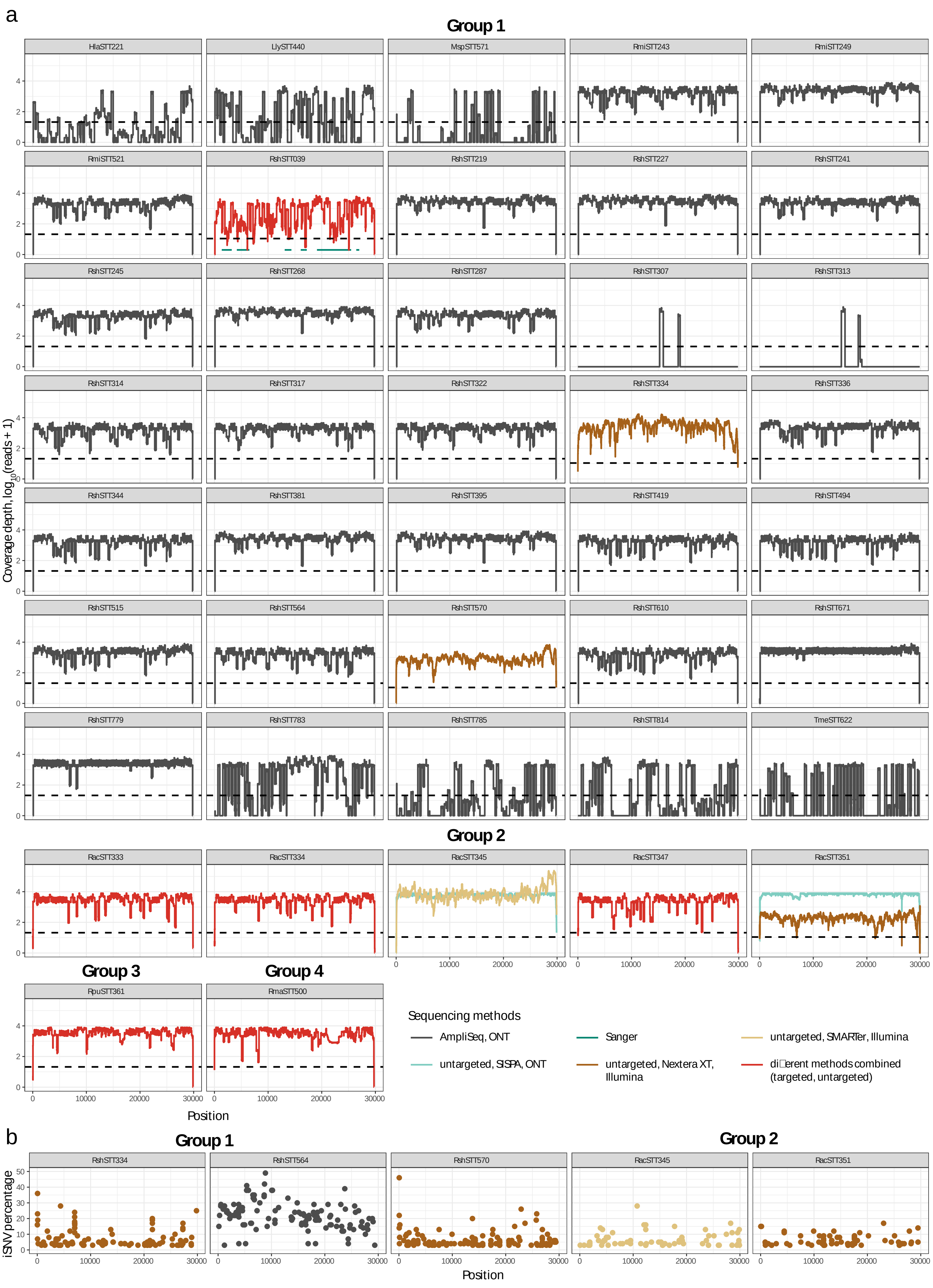

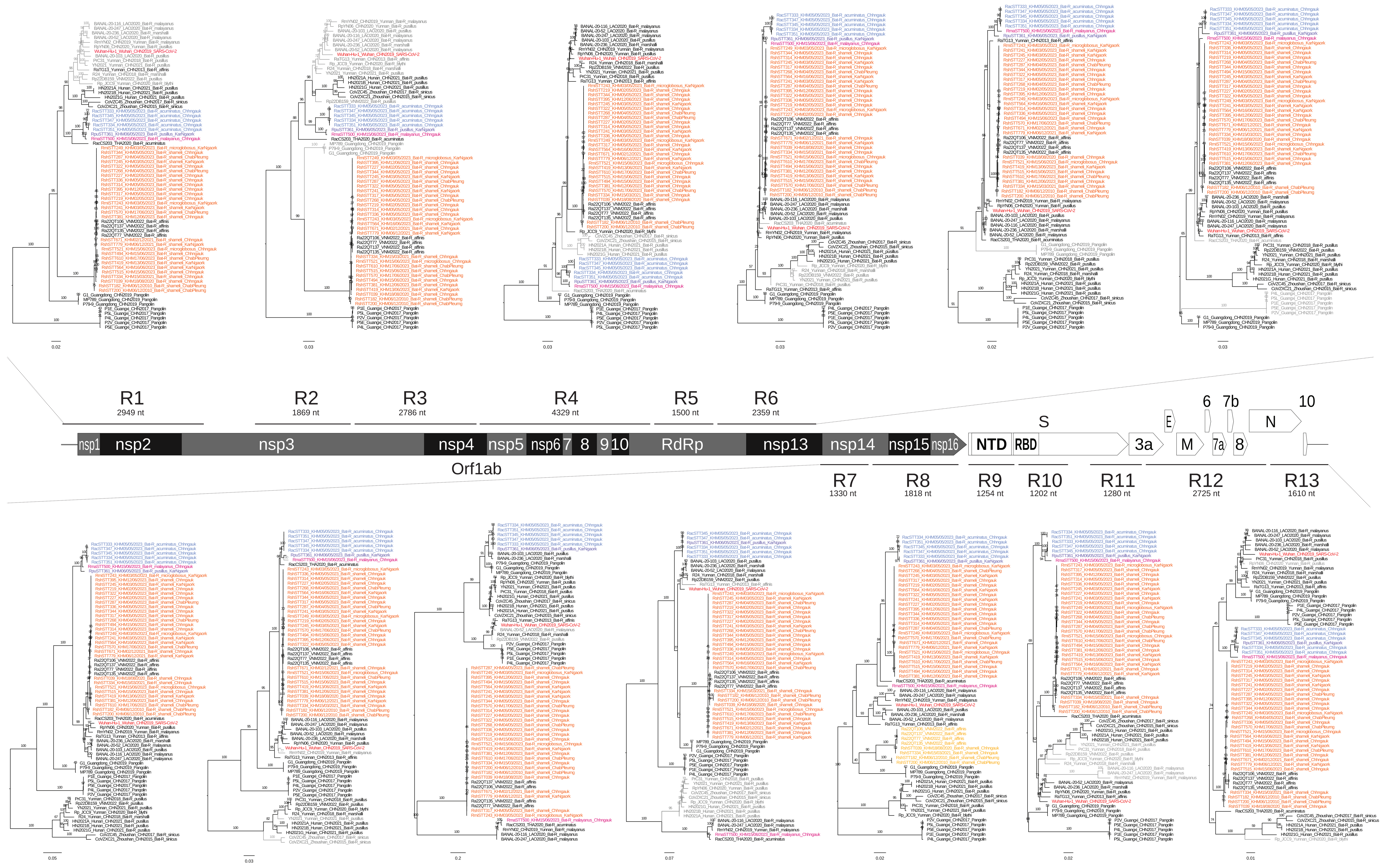

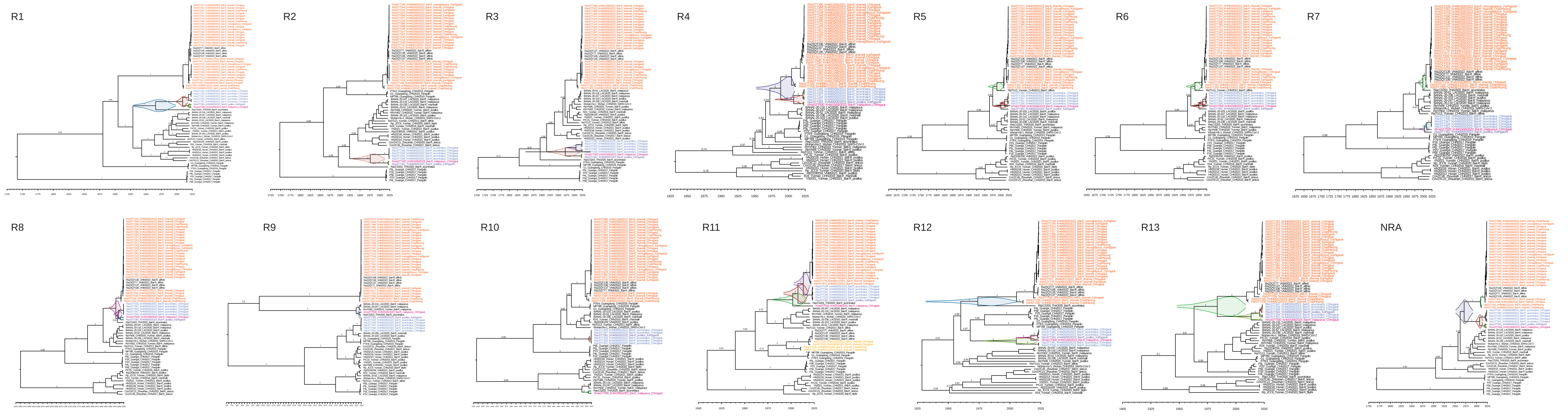

Supplementary Figure 4

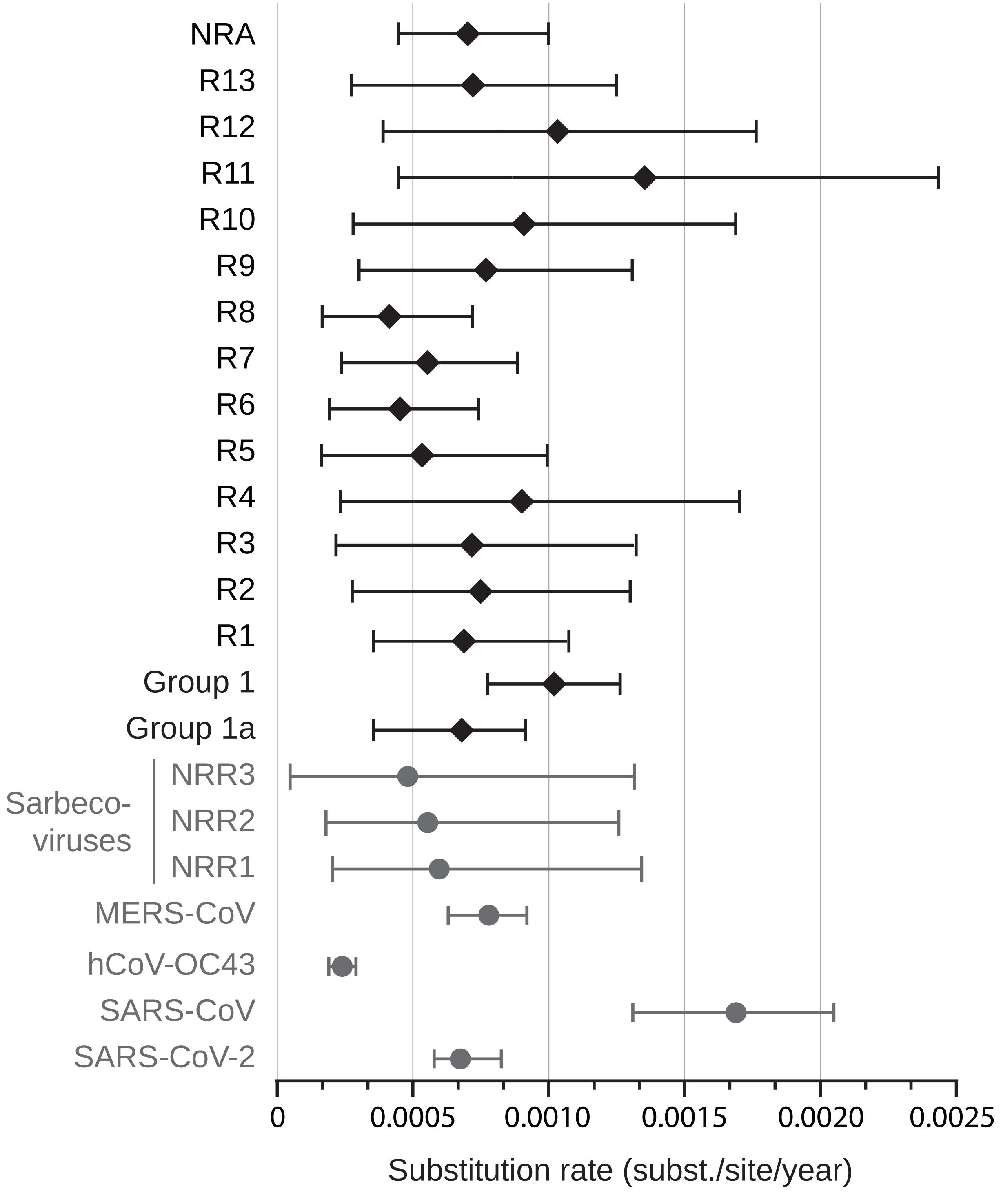

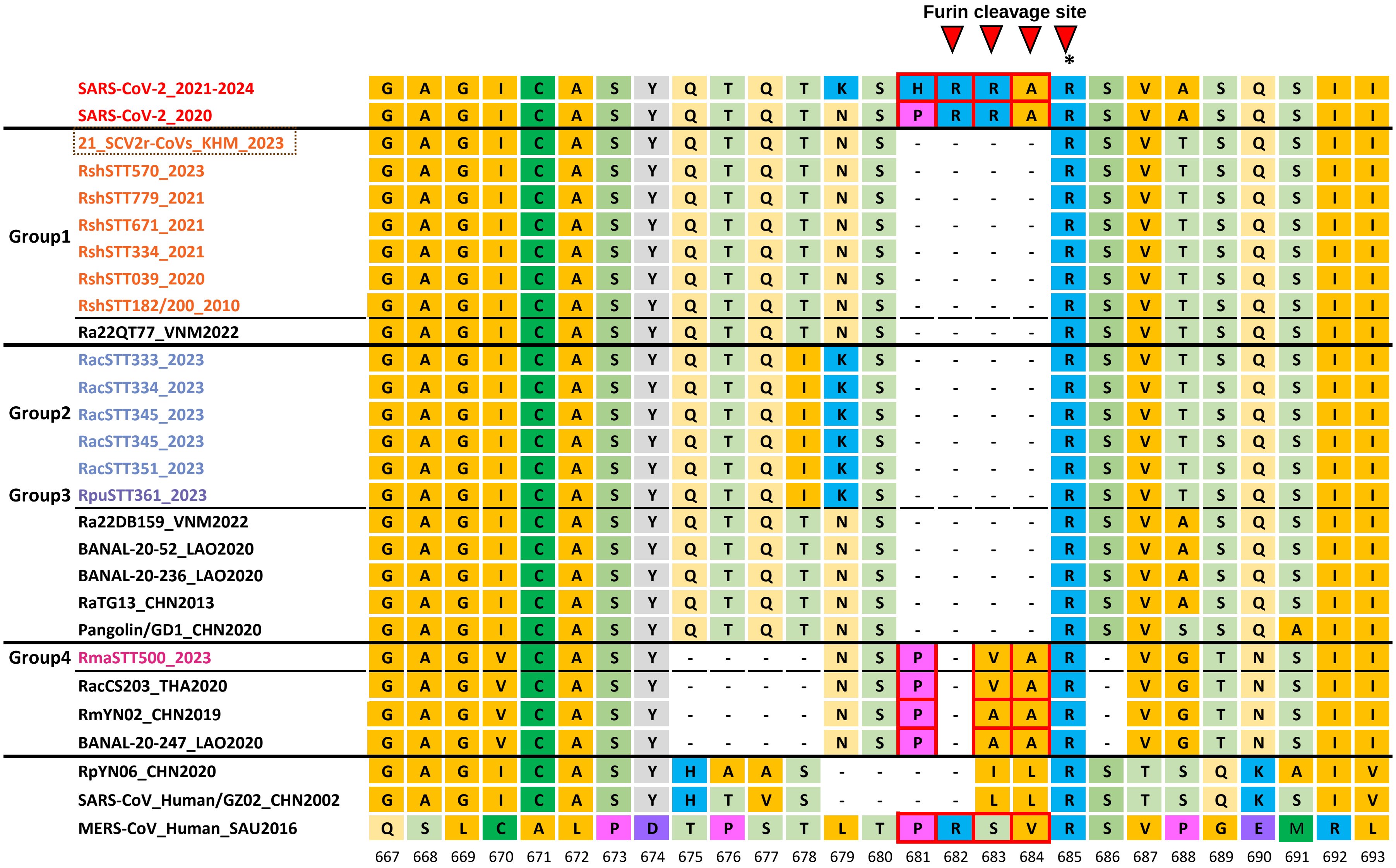
