## Extended_Data_Table for "Characterization and evolutionary history of novel SARS-CoV-2-related viruses in bats from Cambodia"

**Extended Data Table 1:** Bat species captured by date of collection and tested by sarbecovirus real-time RT-qPCR^19^ and pan-coronavirus RT-PCRs^20,21^, at the four sites during active, longitudinal surveillance in Steung Treng, Cambodia, between 2020 and 2023

| **Genus/Species** | **Nb. of samples tested** | **Prevalence of samples positive for CoV (%) by date of sample collection** | | | | | | | | **Nb. of samples positive by sarbecovirus RT-qPCR^a^** | **Nb. of samples positive by pan-CoV RT-PCRs^b^** | **Nb. of samples positive by both RT-PCRs^c^** | **Nb. of samples for sequencing** |
| --- | --- | --- | --- | --- | --- | --- | --- | --- | --- | --- | --- | --- | --- |
|  |  | **Aug-20** | **Oct-20** | **Mar-21** | **Aug-Sep-21** | **Nov-Dec-21** | **Mar-23** | **May-23** | **Jun-23** |  |  |  |  |
| ***Cynopterus spp.*** | 4 |  |  |  | 0/4  (0%) |  |  |  |  |  |  |  |  |
| ***Cynopterus sphinx*** | 28 |  | 0/1 (0%) | 0/6  (0%) | 0/1  (0%) | 0/16 (0%) |  | 0/2  (0%) | 0/2  (0%) |  |  |  |  |
| ***Eonycteris spelaea*** | 6 |  |  | 0/1  (0%) | 0/2  (0%) | 0/1 (0%) |  |  | 0/2  (0%) |  |  |  |  |
| ***Hipposideros armiger*** | 33 | 0/3 (0%) | 0/3 (0%) |  | 0/10 (0%) |  | 0/2  (0%) | **1/9 (11%)** | 0/6  (0%) | **1/33**  **(3%)** |  |  | 1 |
| ***Hipposideros cineraceus*** | 2 |  |  |  |  | 0/2 (0%) |  |  |  |  |  |  |  |
| ***Hipposideros diadema*** | 7 |  |  |  |  |  | 0/1  (0%) | 0/5  (0%) | **1/1 (100%)** |  | **1/7**  **(14%)** |  |  |
| ***Hipposideros galeritus*** | 5 |  | 0/1 (0%) | 0/1  (0%) |  | 0/1 (0%) | 0/1  (0%) | 0/1  (0%) |  |  |  |  |  |
| ***Hipposideros larvatus*** | 55 | **3/13 (23%)** | 0/4 (0%) | 0/3  (0%) | 0/2  (0%) | 0/1 (0%) | 0/3  (0%) | **1/13 (8%)** | **8/16 (50%)** | **1/55**  **(2%)** | **11/55 (20%)** |  | 1 |
| ***Hipposideros pomona*** | 11 |  | **1/3 (33%)** |  |  |  | **1/2 (50%)** | 0/1  (0%) | **3/5**  **(60%)** |  | **5/11**  **(45%)** |  |  |
| ***Hypsugo dolichodon*** | 2 |  |  |  |  |  |  |  | 0/2  (0%) |  |  |  |  |
| ***Lyroderma lyra*** | 39 | 0/4 (0%) | 0/7 (0%) |  | 0/3  (0%) |  |  | 0/2  (0%) | **1/23 (4%)** | **1/39**  **(3%)** |  |  | 1 |
| ***Megaderma spasma*** | 30 | 0/2 (0%) | 0/5 (0%) | 0/2  (0%) | 0/1  (0%) | **1/5 (20%)** | 0/4  (0%) | 0/4  (0%) | **1/7 (14%)** | **1/30**  **(3%)** | **1/30**  **(3%)** |  | 1 |
| ***Rhinolophus acuminatus*** | 8 |  |  |  |  |  |  | **5/8 (63%)** |  |  |  | **5/8**  **(63%)** | 5 |
| ***Rhinolophus chaseni*** | 14 |  |  |  |  | 0/2 (0%) | 0/1  (0%) | **3/10 (30%)** | 0/1  (0%) | **2/14**  **(14%)** | **1/14**  **(7%)** |  | 2 |
| ***Rhinolophus malayanus*** | 40 | 0/2 (0%) | 0/15 (0%) | 0/10 (0%) |  | 0/1 (0%) | 0/7  (0%) | 0/1  (0%) | **1/4 (25%)** |  |  | **1/40**  **(3%)** | 1 |
| ***Rhinolophus microglobosus*** | 34 | **2/7 (29%)** | **1/4 (25%)** | 0/2  (0%) |  | **3/9 (33%)** | 0/6  (0%) | **3/3 (100%)** | **2/3 (67%)** | **1/34**  **(3%)** | **7/34**  **(21%)** | **3/34**  **(9%)** | 4 |
| ***Rhinolophus pusillus*** | 14 |  |  |  |  | 0/5 (0%) | 0/3  (0%) | **1/2 (50%)** | 0/4  (0%) |  |  | **1/14**  **(7%)** | 1 |
| ***Rhinolophus shameli*** | 872 | **5/123 (4%)** | **4/80 (5%)** | **2/75 (3%)** | **1/63 (2%)** | **5/165 (3%)** | **10/158 (6%)** | **41/92 (45%)** | **33/116 (28%)** | **15/872**  **(2%)** | **29/872 (3%)** | **57/872 (7%)** | 32# |
| ***Rhinolophus spp.*** | 5 |  |  |  |  | 0/5 (0%) |  |  |  |  |  |  |  |
| ***Rousettus amplexicaudatus*** | 11 |  |  |  |  | 0/11 (0%) |  |  |  |  |  |  |  |
| ***Rousettus spp.*** | 5 |  |  |  |  | 0/5 (0%) |  |  |  |  |  |  |  |
| ***Taphozous melanopogon*** | 237 | 0/11 (0%) | 0/15 (0%) | 0/17 (0%) | 0/40 (0%) | **1/92 (1%)** | 0/14 (0%) | 0/11 (0%) | **1/37**  **(3%)** | **1/237**  **(0.4%)** | **1/237 (0.4%)** |  | 1 |
| **Total** | 1462 | **10/165 (6%)** | **6/138 (4%)** | **2/117 (2%)** | **1/126 (0.8%)** | **10/321 (3%)** | **11/202 (5%)** | **55/164 (34%)** | **51/229 (22%)** | **23/1462**  **(2%)** | **56/1462 (4%)** | **67/1462 (5%)** | 50 |

Nb.: Number, CoV: coronavirus

^a^Real-time RT-qPCR detecting sarbecoviruses using a duplex assay targeting the E and N genes (19)

^b^RT- PCRs detecting pan-coronaviruses targeting the RdRp gene (20,21)

^c^One bat sample was positive for both the sarbecovirus real-time RT-qPCR and the pan-coronaviruses RT-PCRs.

\# Positive *Rhinolophus shameli* bat samples in 2023 were selected for sequencing based on a Ct value lower than 31, as well as their geographic locations and sampling dates

Bold values indicate a prevalence greater than 0%.
