## Supplementary_Tables for "Characterization and evolutionary history of novel SARS-CoV-2-related viruses in bats from Cambodia"

**Supplementary Table S1. Summary of 146 positive and 29 negative bat samples tested by Sarbecovirus RT-qPCR and Pan-coronaviruses RT-PCRs used for species identification and ACE2 sequencing**

| Sample ID | Genus/Species | Site of sample collection | Date of sampling | Sarbecovirus RT-qPCR (19) | | Pan-CoV RT-PCR | | ID GenBank of partial RdRp gene | Classification based on partial RdRp sequence | COI GenBank ID | ACE2 GenBank ID |
|---|---|---|---|---|---|---|---|---|---|---|---|
| | | | | E-gene Ct value | N-gene Ct value | Quan et al (20) | Chu et al (21) | | | | |
| RshSTT039 | *Rhinolophus shameli* | Phnom Chhngauk | 2020-08-18 | 29.22 | 27.33 | Negative | NA | PQ305308 | BetaCoV-Sarbecovirus | OZ248192 | OZ248119 |
| RshSTT334 | *Rhinolophus shameli* | Phnom Chhngauk | 2021-03-15 | 24.35 | 23.5 | Negative | NA | PQ305309 | BetaCoV-Sarbecovirus | OZ248193 | NA |
| MspSTT571 | *Megaderma spasma* | Phnom Chab Phleurng | 2021-11-30 | 38.4 | 38.27 | Negative | NA | PQ305310 | BetaCoV-Sarbecovirus | OZ248194 | OZ248090 |
| TmeSTT622 | *Taphozous melanopogon* | Phnom Ka Ngoak | 2021-12-01 | 37.99 | 36.38 | Negative | NA | PQ305311 | BetaCoV-Sarbecovirus | OZ248195 | OZ248094 |
| RshSTT671 | *Rhinolophus shameli* | Phnom Chhngauk | 2021-12-02 | 21.22 | 19.21 | Positive | NA | PQ305312 | BetaCoV-Sarbecovirus | OZ248196 | OZ248124 |
| RshSTT779 | *Rhinolophus shameli* | Phnom Ka Ngoak | 2021-12-06 | 23.98 | 22.23 | Negative | NA | PQ305313 | BetaCoV-Sarbecovirus | OZ248197 | OZ248125 |
| RshSTT783 | *Rhinolophus shameli* | Phnom Ka Ngoak | 2021-12-06 | 34.38 | 33.34 | Negative | NA | PQ305314 | BetaCoV-Sarbecovirus | OZ248198 | OZ248126 |
| RshSTT785 | *Rhinolophus shameli* | Phnom Ka Ngoak | 2021-12-06 | 40.63 | 36.77 | Negative | NA | NA | BetaCoV-Sarbecovirus | OZ248199 | OZ248127 |
| RshSTT814 | *Rhinolophus shameli* | Phnom Ka Ngoak | 2021-12-06 | 36.77 | 37.03 | Negative | NA | PQ305315 | BetaCoV-Sarbecovirus | OZ248200 | OZ248128 |
| RshSTT003 | *Rhinolophus shameli* | Phnom Chhngauk | 2020-08-18 | Negative | Negative | Positive | NA | PQ305323 | AlphaCoV-Decacovirus | OZ248136 | OZ248116 |
| RmiSTT018 | *Rhinolophus microglobosus* | Phnom Chhngauk | 2020-08-18 | Negative | Negative | Positive | NA | PQ305324 | AlphaCoV-Rhinacovirus | OZ248201 | OZ248104 |
| RshSTT025 | *Rhinolophus shameli* | Phnom Chhngauk | 2020-08-18 | Negative | Negative | Positive | NA | PQ305325 | AlphaCoV-Decacovirus | OZ248137 | OZ248117 |
| RshSTT028 | *Rhinolophus shameli* | Phnom Chhngauk | 2020-08-18 | Negative | Negative | Positive | NA | PQ305326 | AlphaCoV-Rhinacovirus | OZ248138 | OZ248118 |
| HlaSTT087 | *Hipposideros larvatus* | Phnom Kork Romeas + Chab Phleurng | 2020-08-19 | Negative | Negative | Positive | NA | PQ305327 | AlphaCoV-Decacovirus | OZ248202 | NA |
| RshSTT096 | *Rhinolophus shameli* | Phnom Kork Romeas + Chab Phleurng | 2020-08-19 | Negative | Negative | Positive | NA | PQ305328 | AlphaCoV-Decacovirus | OZ248203 | NA |
| HlaSTT101 | *Hipposideros larvatus* | Phnom Kork Romeas + Chab Phleurng | 2020-08-19 | Negative | Negative | Positive | NA | PQ305329 | AlphaCoV-Decacovirus | OZ248204 | NA |
| HlaSTT106 | *Hipposideros larvatus* | Phnom Chhngauk | 2020-08-20 | Negative | Negative | Positive | NA | PQ305330 | AlphaCoV-Decacovirus | OZ248205 | NA |
| RmiSTT108 | *Rhinolophus microglobosus* | Phnom Chhngauk | 2020-08-20 | Negative | Negative | Positive | NA | PQ305331 | AlphaCoV-Decacovirus | OZ248139 | NA |
| RshSTT175 | *Rhinolophus shameli* | Phnom Chhngauk | 2020-10-13 | Negative | Negative | Positive | NA | PQ305332 | AlphaCoV-Decacovirus | OZ248140 | OZ248120 |
| RmiSTT211 | *Rhinolophus microglobosus* | Phnom Chhngauk | 2020-10-15 | Negative | Negative | Positive | NA | PQ305333 | AlphaCoV-Rhinacovirus | OZ248206 | NA |
| RshSTT221 | *Rhinolophus shameli* | Phnom Chhngauk | 2020-10-15 | Negative | Negative | Positive | NA | PQ305334 | AlphaCoV-Decacovirus | OZ248141 | NA |
| HpoSTT222 | *Hipposideros pomona* | Phnom Chhngauk | 2020-10-15 | Negative | Negative | Positive | NA | PQ305335 | AlphaCoV-Decacovirus | OZ248142 | OZ248084 |
| RshSTT279 | *Rhinolophus shameli* | Phnom Ka Ngoak | 2020-10-16 | Negative | Negative | Positive | NA | PQ305336 | AlphaCoV-Rhinacovirus | OZ248143 | OZ248121 |
| RshSTT300 | *Rhinolophus shameli* | Phnom Chhngauk | 2020-10-17 | Negative | Negative | Positive | NA | PQ305337 | AlphaCoV-Decacovirus | OZ248144 | OZ248122 |
| RshSTT404 | *Rhinolophus shameli* | Phnom Ka Ngoak | 2021-03-17 | Negative | Negative | Positive | NA | PQ305316 | AlphaCoV-Decacovirus | OZ248145 | OZ248123 |
| RshSTT543 | *Rhinolophus shameli* | Phnom Kork Romeas | 2021-09-03 | Negative | Negative | Positive | NA | PQ305317 | AlphaCoV-Decacovirus | OZ248207 | NA |
| RmiSTT589 | *Rhinolophus microglobosus* | Phnom Ka Ngoak | 2021-12-01 | Negative | Negative | Positive | NA | PQ305318 | AlphaCoV-Rhinacovirus | OZ248208 | NA |
| RmiSTT611 | *Rhinolophus microglobosus* | Phnom Ka Ngoak | 2021-12-01 | Negative | Negative | Positive | NA | PQ305338 | AlphaCoV-Rhinacovirus | OZ248146 | OZ248107 |
| RmiSTT794 | *Rhinolophus microglobosus* | Phnom Ka Ngoak | 2021-12-06 | Negative | Negative | Positive | NA | PQ305319 | AlphaCoV-Rhinacovirus | OZ248209 | OZ248109 |
| HlaSTT009 | *Hipposideros larvatus* | Phnom Chhngauk | 2020-08-18 | Negative | Negative | Negative | NA | NA | NA | OZ248147 | OZ248078 |
| MspSTT013 | *Megaderma spasma* | Phnom Chhngauk | 2020-08-18 | Negative | Negative | Negative | NA | NA | NA | OZ248148 | OZ248088 |
| HlaSTT019 | *Hipposideros larvatus* | Phnom Chhngauk | 2020-08-18 | Negative | Negative | Negative | NA | NA | NA | OZ248149 | OZ248079 |
| RmaSTT060 | *Rhinolophus malayanus* | Phnom Kork Romeas | 2020-08-19 | Negative | Negative | Negative | NA | NA | NA | OZ248150 | OZ248097 |
| TmeSTT071 | *Taphozous melanopogon* | Phnom Kork Romeas | 2020-08-19 | Negative | Negative | Negative | NA | NA | NA | OZ248151 | OZ248091 |
| HlaSTT107 | *Hipposideros larvatus* | Phnom Chhngauk | 2020-08-20 | Negative | Negative | Negative | NA | NA | NA | OZ248152 | OZ248080 |
| LlySTT109 | *Lyroderma lyra* | Phnom Chhngauk | 2020-08-20 | Negative | Negative | Negative | NA | NA | NA | OZ248153 | OZ248085 |
| HarSTT118 | *Hipposideros armiger* | Phnom Ka Ngoak | 2020-08-21 | Negative | Negative | Negative | NA | NA | NA | OZ248154 | OZ248075 |
| HlaSTT183 | *Hipposideros larvatus* | Phnom Chhngauk | 2020-10-13 | Negative | Negative | Negative | NA | NA | NA | OZ248155 | OZ248081 |
| RmaSTT187 | *Rhinolophus malayanus* | Phnom Chhngauk | 2020-10-13 | Negative | Negative | Negative | NA | NA | NA | OZ248156 | OZ248098 |
| LlySTT189 | *Lyroderma lyra* | Phnom Chhngauk | 2020-10-13 | Negative | Negative | Negative | NA | NA | NA | OZ248157 | OZ248086 |
| HpoSTT218 | *Hipposideros pomona* | Phnom Chhngauk | 2020-10-15 | Negative | Negative | Negative | NA | NA | NA | OZ248158 | OZ248083 |
| MspSTT236 | *Megaderma spasma* | Phnom Chhngauk | 2020-10-15 | Negative | Negative | Negative | NA | NA | NA | OZ248159 | OZ248089 |
| TmeSTT355 | *Taphozous melanopogon* | Phnom Chhngauk | 2021-03-15 | Negative | Negative | Negative | NA | NA | NA | OZ248160 | OZ248092 |
| RmiSTT406 | *Rhinolophus microglobosus* | Phnom Ka Ngoak | 2021-03-17 | Negative | Negative | Negative | NA | NA | NA | OZ248161 | OZ248105 |
| RmiSTT409 | *Rhinolophus microglobosus* | Phnom Ka Ngoak | 2021-03-17 | Negative | Negative | Negative | NA | NA | NA | OZ248162 | OZ248106 |
| TmeSTT436 | *Taphozous melanopogon* | Phnom Chhngauk | 2021-08-31 | Negative | Negative | Negative | NA | NA | NA | OZ248163 | OZ248093 |
| CspSTT575 | *Cynopterus sphinx* | Phnom Chab Phleurng | 2021-11-30 | Negative | Negative | Negative | NA | NA | NA | OZ248164 | OZ248074 |
| CspSTT582 | *Cynopterus sphinx* | Phnom Chab Phleurng | 2021-11-30 | Negative | Negative | Negative | NA | NA | NA | OZ248165 | OZ248073 |
| RmiSTT651 | *Rhinolophus microglobosus* | Phnom Chhngauk | 2021-12-02 | Negative | Negative | Negative | NA | NA | NA | OZ248166 | OZ248108 |
| HgaSTT665 | *Hipposideros galeritus* | Phnom Chhngauk | 2021-12-02 | Negative | Negative | Negative | NA | NA | NA | OZ248167 | OZ248077 |
| RchSTT675 | *Rhinolophus chaseni* | Phnom Kork Romeas | 2021-12-04 | Negative | Negative | Negative | NA | NA | NA | OZ248168 | OZ248095 |
| RchSTT748 | *Rhinolophus chaseni* | Phnom Chab Phleurng | 2021-12-05 | Negative | Negative | Negative | NA | NA | NA | OZ248169 | OZ248096 |
| RmaSTT809 | *Rhinolophus malayanus* | Phnom Ka Ngoak | 2021-12-06 | Negative | Negative | Negative | NA | NA | NA | OZ248170 | OZ248099 |
| RmiSTT810 | *Rhinolophus microglobosus* | Phnom Ka Ngoak | 2021-12-06 | Negative | Negative | Negative | NA | NA | NA | OZ248171 | OZ248110 |
| RmiSTT820 | *Rhinolophus microglobosus* | Phnom Ka Ngoak | 2021-12-06 | Negative | Negative | Negative | NA | NA | NA | OZ248172 | OZ248111 |
| RmiSTT821 | *Rhinolophus microglobosus* | Phnom Ka Ngoak | 2021-12-06 | Negative | Negative | Negative | NA | NA | NA | OZ248173 | OZ248112 |
| RmiSTT832 | *Rhinolophus microglobosus* | Phnom Ka Ngoak | 2021-12-06 | Negative | Negative | Negative | NA | NA | NA | OZ248174 | OZ248113 |
| RmiSTT846 | *Rhinolophus microglobosus* | Phnom Ka Ngoak | 2021-12-06 | Negative | Negative | Negative | NA | NA | NA | OZ248175 | OZ248114 |
| RshSTT219 | *Rhinolophus shameli* | Phnom Chhngauk | 2023-05-02 | 24.29 | 24.02 | Positive | Positive | OZ250989/OZ250910 | BetaCoV-Sarbecovirus | OZ248210 | NA |
| HlaSTT221 | *Hipposideros larvatus* | Phnom Chhngauk | 2023-05-02 | 39.25 | 38.74 | Negative | Negative | NA | BetaCoV-Sarbecovirus | OZ248211 | OZ248082 |
| RshSTT227 | *Rhinolophus shameli* | Phnom Chhngauk | 2023-05-02 | 23.29 | 23.17 | Positive | Positive | OZ250990/OZ250911 | BetaCoV-Sarbecovirus | OZ248212 | NA |
| RshSTT241 | *Rhinolophus shameli* | Phnom Ka Ngoak | 2023-05-03 | 28.08 | 27.05 | Positive | Positive | OZ250992/OZ250913 | BetaCoV-Sarbecovirus | OZ248213 | NA |
| RmiSTT243 | *Rhinolophus microglobosus* | Phnom Ka Ngoak | 2023-05-03 | 17.86 | 15.76 | Positive | Positive | OZ250993/OZ250914 | BetaCoV-Sarbecovirus | OZ248176 | NA |
| RshSTT245 | *Rhinolophus shameli* | Phnom Ka Ngoak | 2023-05-03 | 17.05 | 16.86 | Positive | Positive | OZ250994/OZ250915 | BetaCoV-Sarbecovirus | OZ248214 | NA |
| RshSTT246 | *Rhinolophus shameli* | Phnom Ka Ngoak | 2023-05-03 | 35.9 | 36.05 | Negative | Positive | OZ250884 | BetaCoV-Sarbecovirus | OZ248215 | NA |
| RmiSTT249 | *Rhinolophus microglobosus* | Phnom Ka Ngoak | 2023-05-03 | 23.04 | 22.06 | Positive | Positive | OZ250995/OZ250916 | BetaCoV-Sarbecovirus | OZ248177 | NA |
| RmiSTT252 | *Rhinolophus microglobosus* | Phnom Ka Ngoak | 2023-05-03 | 39.79 | 37.47 | Negative | Negative | NA | Sarbecovirus positive by RT-qPCR | OZ248178 | NA |
| RshSTT260 | *Rhinolophus shameli* | Phnom Ka Ngoak | 2023-05-03 | 27.17 | 25.98 | Positive | Positive | OZ250996/OZ250917 | BetaCoV-Sarbecovirus | OZ248216 | NA |
| RshSTT265 | *Rhinolophus shameli* | Phnom Chab Phleurng | 2023-05-04 | 37.47 | 37.68 | Negative | Negative | NA | Sarbecovirus positive by RT-qPCR | OZ248217 | NA |
| RshSTT268 | *Rhinolophus shameli* | Phnom Chab Phleurng | 2023-05-04 | 21.46 | 21.12 | Positive | Positive | OZ250997/OZ250918 | BetaCoV-Sarbecovirus | OZ248218 | NA |
| RshSTT271 | *Rhinolophus shameli* | Phnom Chab Phleurng | 2023-05-04 | 22.39 | 22.2 | Positive | Positive | OZ250998/OZ250919 | BetaCoV-Sarbecovirus | OZ248219 | NA |
| RshSTT273 | *Rhinolophus shameli* | Phnom Chab Phleurng | 2023-05-04 | 28.49 | 26.81 | Negative | Positive | OZ250885 | BetaCoV-Sarbecovirus | OZ248220 | NA |
| RshSTT280 | *Rhinolophus shameli* | Phnom Chab Phleurng | 2023-05-04 | 25.29 | 24.13 | Positive | Positive | OZ251001/OZ250922 | BetaCoV-Sarbecovirus | OZ248221 | NA |
| RshSTT284 | *Rhinolophus shameli* | Phnom Chab Phleurng | 2023-05-04 | 34.09 | 33.84 | Positive | Positive | OZ251002/OZ250923 | BetaCoV-Sarbecovirus | OZ248222 | NA |
| RshSTT286 | *Rhinolophus shameli* | Phnom Chab Phleurng | 2023-05-04 | 24.03 | 22.16 | Positive | Positive | OZ251003/OZ250924 | BetaCoV-Sarbecovirus | OZ248223 | NA |
| RshSTT287 | *Rhinolophus shameli* | Phnom Chab Phleurng | 2023-05-04 | 23.5 | 21.45 | Positive | Positive | OZ251004/OZ250925 | BetaCoV-Sarbecovirus | OZ248224 | NA |
| RshSTT288 | *Rhinolophus shameli* | Phnom Chab Phleurng | 2023-05-04 | 37.12 | 36.22 | Positive | Positive | OZ251005/OZ250926 | BetaCoV-sarbecovirus & AlphaCoV-Decacovirus | OZ248225 | NA |
| RshSTT290 | *Rhinolophus shameli* | Phnom Chab Phleurng | 2023-05-04 | 38.61 | 37.65 | Negative | Negative | NA | Sarbecovirus positive by RT-qPCR | OZ248226 | NA |
| RshSTT292 | *Rhinolophus shameli* | Phnom Chab Phleurng | 2023-05-04 | 31.37 | 30.65 | Positive | Positive | OZ251006/OZ250927 | BetaCoV-Sarbecovirus | OZ248227 | NA |
| RshSTT293 | *Rhinolophus shameli* | Phnom Chab Phleurng | 2023-05-04 | 28.68 | 29.46 | Positive | Positive | OZ251007/OZ250928 | BetaCoV-Sarbecovirus | OZ248228 | NA |
| RshSTT295 | *Rhinolophus shameli* | Phnom Chab Phleurng | 2023-05-04 | 32.29 | 31.77 | Negative | Positive | OZ250887 | BetaCoV-Sarbecovirus | OZ248229 | NA |
| RshSTT296 | *Rhinolophus shameli* | Phnom Chab Phleurng | 2023-05-04 | 40.73 | 36.29 | Negative | Negative | NA | Sarbecovirus positive by RT-qPCR | OZ248230 | NA |
| RshSTT298 | *Rhinolophus shameli* | Phnom Chab Phleurng | 2023-05-04 | 34.28 | 34.18 | Positive | Positive | OZ251008/OZ250929 | BetaCoV-Sarbecovirus | OZ248231 | NA |
| RshSTT299 | *Rhinolophus shameli* | Phnom Chab Phleurng | 2023-05-04 | 37.65 | 35.81 | Negative | Positive | OZ250888 | BetaCoV-Sarbecovirus | OZ248232 | NA |
| RshSTT302 | *Rhinolophus shameli* | Phnom Chab Phleurng | 2023-05-04 | 32.64 | 30.3 | Negative | Positive | OZ250889 | BetaCoV-Sarbecovirus | OZ248233 | NA |
| RshSTT307 | *Rhinolophus shameli* | Phnom Chab Phleurng | 2023-05-04 | 36.13 | 33.92 | Positive | Positive | OZ251010/OZ250931 | BetaCoV-sarbecovirus & AlphaCoV-Decacovirus | OZ248234 | NA |
| RshSTT312 | *Rhinolophus shameli* | Phnom Chhngauk | 2023-05-05 | 32.94 | 34.57 | Positive | Positive | OZ251011/OZ250932 | BetaCoV-Sarbecovirus | OZ248235 | NA |
| RshSTT313 | *Rhinolophus shameli* | Phnom Chhngauk | 2023-05-05 | 36.65 | 36.33 | Positive | Positive | OZ251012/OZ250933 | BetaCoV-sarbecovirus & AlphaCoV-Decacovirus | OZ248236 | NA |
| RshSTT314 | *Rhinolophus shameli* | Phnom Chhngauk | 2023-05-05 | 20.87 | 18.09 | Positive | Positive | OZ251013/OZ250934 | BetaCoV-Sarbecovirus | OZ248237 | NA |
| RshSTT316 | *Rhinolophus shameli* | Phnom Chhngauk | 2023-05-05 | 31.84 | 30.75 | Positive | Positive | OZ251014/OZ250935 | BetaCoV-Sarbecovirus | OZ248238 | NA |
| RshSTT317 | *Rhinolophus shameli* | Phnom Chhngauk | 2023-05-05 | 20.15 | 17.34 | Positive | Positive | OZ251015/OZ250936 | BetaCoV-Sarbecovirus | OZ248239 | NA |
| RshSTT322 | *Rhinolophus shameli* | Phnom Chhngauk | 2023-05-05 | 21.32 | 19.14 | Positive | Positive | OZ251016/OZ250937 | BetaCoV-Sarbecovirus | OZ248240 | NA |
| RshSTT327 | *Rhinolophus shameli* | Phnom Chhngauk | 2023-05-05 | 33.37 | 31.43 | Negative | Positive | OZ250890 | BetaCoV-Sarbecovirus | OZ248241 | NA |
| RshSTT328 | *Rhinolophus shameli* | Phnom Chhngauk | 2023-05-05 | 25.02 | 22.61 | Positive | Positive | OZ251018/OZ250939 | BetaCoV-Sarbecovirus | OZ248242 | NA |
| RshSTT331 | *Rhinolophus shameli* | Phnom Chhngauk | 2023-05-05 | 25.41 | 23.4 | Positive | Positive | OZ251019/OZ250940 | BetaCoV-Sarbecovirus | OZ248243 | NA |
| RacSTT333 | *Rhinolophus acuminatus* | Phnom Chhngauk | 2023-05-05 | 29.07 | 28.72 | Positive | Positive | OZ251020/OZ250941 | BetaCoV-Sarbecovirus | OZ248244 | NA |
| RacSTT334 | *Rhinolophus acuminatus* | Phnom Chhngauk | 2023-05-05 | 32.12 | 31.42 | Positive | Positive | OZ251021/OZ250942 | BetaCoV-Sarbecovirus | OZ248179 | OZ248101 |
| RshSTT336 | *Rhinolophus shameli* | Phnom Chhngauk | 2023-05-05 | 19.27 | 17.78 | Positive | Positive | OZ251022/OZ250943 | BetaCoV-Sarbecovirus | OZ248245 | NA |
| RchSTT337 | *Rhinolophus chaseni* | Phnom Chhngauk | 2023-05-05 | 39.58 | 38.37 | Negative | Negative | NA | Sarbecovirus positive by RT-qPCR | OZ248246 | NA |
| RshSTT344 | *Rhinolophus shameli* | Phnom Chhngauk | 2023-05-05 | 19.05 | 17.33 | Positive | Positive | OZ251023/OZ250944 | BetaCoV-Sarbecovirus | OZ248247 | NA |
| RacSTT345 | *Rhinolophus acuminatus* | Phnom Chhngauk | 2023-05-05 | 17.68 | 15.52 | Positive | Positive | OZ251024/OZ250945 | BetaCoV-Sarbecovirus | OZ248180 | OZ248102 |
| RacSTT347 | *Rhinolophus acuminatus* | Phnom Chhngauk | 2023-05-05 | 30.51 | 28.78 | Positive | Positive | OZ251025/OZ250946 | BetaCoV-Sarbecovirus | OZ248181 | NA |
| RacSTT351 | *Rhinolophus acuminatus* | Phnom Chhngauk | 2023-05-05 | 22.13 | 19.66 | Positive | Positive | OZ251026/OZ250947 | BetaCoV-Sarbecovirus | OZ248182 | NA |
| RchSTT352 | *Rhinolophus chaseni* | Phnom Chhngauk | 2023-05-05 | 38.19 | 37.25 | Negative | Negative | NA | Sarbecovirus positive by RT-qPCR | OZ248183 | NA |
| RpuSTT361 | *Rhinolophus pusillus* | Phnom Ka Ngoak | 2023-05-06 | 31.41 | 33.09 | Positive | Positive | OZ251027/OZ250948 | BetaCoV-Sarbecovirus | OZ248248 | OZ248103 |
| HarSTT369 | *Hipposideros armiger* | Phnom Ka Ngoak | 2023-05-06 | 38.9 | 38.47 | Negative | Negative | NA | Sarbecovirus positive by RT-qPCR | OZ248249 | OZ248076 |
| RshSTT381 | *Rhinolophus shameli* | Phnom Chhngauk | 2023-06-12 | 23.66 | 20.39 | Positive | Positive | OZ251028/OZ250949 | BetaCoV-Sarbecovirus | OZ248250 | NA |
| RshSTT395 | *Rhinolophus shameli* | Phnom Chhngauk | 2023-06-12 | 24.85 | 23.64 | Positive | Positive | OZ251032/OZ250953 | BetaCoV-Sarbecovirus | OZ248251 | NA |
| RshSTT419 | *Rhinolophus shameli* | Phnom Ka Ngoak | 2023-06-13 | 19.33 | 18.45 | Negative | Positive | OZ250892 | BetaCoV-Sarbecovirus | OZ248252 | NA |
| LlySTT440 | *Lyroderma lyra* | Phnom Ka Ngoak | 2023-06-13 | 37.94 | 36.87 | Negative | Negative | NA | BetaCoV-Sarbecovirus | OZ248184 | OZ248087 |
| RshSTT463 | *Rhinolophus shameli* | Phnom Chab Phleurng | 2023-06-14 | 38.33 | 37.88 | Negative | Negative | NA | Sarbecovirus positive by RT-qPCR | OZ248253 | NA |
| RshSTT470 | *Rhinolophus shameli* | Phnom Chab Phleurng | 2023-06-14 | 36.01 | 35.08 | Negative | Positive | OZ250894 | BetaCoV-Sarbecovirus | OZ248254 | NA |
| RshSTT488 | *Rhinolophus shameli* | Phnom Chab Phleurng | 2023-06-14 | 34.72 | 34.58 | Positive | Negative | OZ250978 | BetaCoV-Sarbecovirus | OZ248255 | NA |
| RshSTT491 | *Rhinolophus shameli* | Phnom Chab Phleurng | 2023-06-14 | 39.27 | 38.35 | Negative | Negative | NA | Sarbecovirus positive by RT-qPCR | OZ248256 | NA |
| RshSTT494 | *Rhinolophus shameli* | Phnom Chhngauk | 2023-06-15 | 21.08 | 19.68 | Positive | Positive | OZ251037/OZ250958 | BetaCoV-Sarbecovirus | OZ248257 | NA |
| RmaSTT500 | *Rhinolophus malayanus* | Phnom Chhngauk | 2023-06-15 | 27.47 | 28.45 | Positive | Positive | OZ251038/OZ250959 | BetaCoV-Sarbecovirus | OZ248185 | OZ248100 |
| RshSTT502 | *Rhinolophus shameli* | Phnom Chhngauk | 2023-06-15 | 26.51 | 24.89 | Positive | Positive | OZ251039/OZ250960 | BetaCoV-Sarbecovirus | OZ248258 | NA |
| RshSTT503 | *Rhinolophus shameli* | Phnom Chhngauk | 2023-06-15 | 34.86 | 34.27 | Negative | Positive | OZ250896 | BetaCoV-sarbecovirus & BetaCoV-Hibecovirus | OZ248259 | NA |
| RshSTT511 | *Rhinolophus shameli* | Phnom Chhngauk | 2023-06-15 | 28.01 | 25.92 | Positive | Negative | OZ250979 | BetaCoV-Sarbecovirus | OZ248260 | NA |
| RshSTT512 | *Rhinolophus shameli* | Phnom Chhngauk | 2023-06-15 | 25.53 | 23.66 | Positive | Positive | OZ251042/OZ250963 | BetaCoV-Sarbecovirus | OZ248261 | NA |
| RshSTT515 | *Rhinolophus shameli* | Phnom Chhngauk | 2023-06-15 | 18.65 | 17.22 | Positive | Positive | OZ251043/OZ250964 | BetaCoV-Sarbecovirus | OZ248262 | NA |
| RshSTT516 | *Rhinolophus shameli* | Phnom Chhngauk | 2023-06-15 | 33.8 | 31.44 | Positive | Positive | OZ251044/OZ250965 | BetaCoV-Sarbecovirus | OZ248263 | NA |
| RshSTT517 | *Rhinolophus shameli* | Phnom Chhngauk | 2023-06-15 | 35.69 | 35.21 | Negative | Positive | OZ250897 | BetaCoV-sarbecovirus & AlphaCoV-Decacovirus | OZ248264 | NA |
| RshSTT518 | *Rhinolophus shameli* | Phnom Chhngauk | 2023-06-15 | 30.93 | 29.43 | Negative | Positive | OZ250898 | BetaCoV-Sarbecovirus | OZ248265 | NA |
| RmiSTT521 | *Rhinolophus microglobosus* | Phnom Chhngauk | 2023-06-15 | 28.29 | 26.39 | Positive | Negative | OZ250980 | BetaCoV-sarbecovirus & AlphaCoV-Rhinacovirus | OZ248186 | OZ248115 |
| RshSTT523 | *Rhinolophus shameli* | Phnom Chhngauk | 2023-06-15 | 38.18 | 39.38 | Negative | Positive | OZ250900 | BetaCoV-sarbecovirus & BetaCoV-Hibecovirus | OZ248266 | NA |
| RshSTT524 | *Rhinolophus shameli* | Phnom Chhngauk | 2023-06-15 | 26.55 | 24.8 | Positive | Positive | OZ251045/OZ250966 | BetaCoV-Sarbecovirus | OZ248267 | NA |
| RshSTT562 | *Rhinolophus shameli* | Phnom Ka Ngoak | 2023-06-16 | 25.17 | 24.02 | Positive | Positive | OZ251047/OZ250968 | BetaCoV-Sarbecovirus | OZ248268 | NA |
| RshSTT564 | *Rhinolophus shameli* | Phnom Ka Ngoak | 2023-06-16 | 21.01 | 19.27 | Positive | Positive | OZ251048/OZ250969 | BetaCoV-Sarbecovirus | OZ248269 | NA |
| RshSTT565 | *Rhinolophus shameli* | Phnom Ka Ngoak | 2023-06-16 | 29.49 | 29.04 | Positive | Positive | OZ251049/OZ250970 | BetaCoV-Sarbecovirus | OZ248270 | NA |
| RshSTT570 | *Rhinolophus shameli* | Phnom Chab Phleurng | 2023-06-17 | 21.55 | 20.28 | Positive | Positive | OZ251050/OZ250971 | BetaCoV-Sarbecovirus | OZ248271 | NA |
| RshSTT575 | *Rhinolophus shameli* | Phnom Chab Phleurng | 2023-06-17 | 33.17 | 32 | Negative | Negative | NA | Sarbecovirus positive by RT-qPCR | OZ248187 | NA |
| RshSTT576 | *Rhinolophus shameli* | Phnom Chab Phleurng | 2023-06-17 | 27.39 | 26.43 | Positive | Positive | OZ251052/OZ250973 | BetaCoV-Sarbecovirus | OZ248272 | NA |
| RshSTT586 | *Rhinolophus shameli* | Phnom Chab Phleurng | 2023-06-17 | 26.36 | 25.3 | Positive | Positive | OZ251053/OZ250974 | BetaCoV-Sarbecovirus | OZ248273 | NA |
| RshSTT587 | *Rhinolophus shameli* | Phnom Chab Phleurng | 2023-06-17 | 33.47 | 32.74 | Negative | Negative | NA | Sarbecovirus positive by RT-qPCR | OZ248188 | NA |
| RshSTT590 | *Rhinolophus shameli* | Phnom Chab Phleurng | 2023-06-17 | 25.15 | 22.28 | Positive | Positive | OZ251054/OZ250975 | BetaCoV-Sarbecovirus | OZ248274 | NA |
| RshSTT599 | *Rhinolophus shameli* | Phnom Chab Phleurng | 2023-06-17 | 39.58 | 39.27 | Negative | Negative | NA | Sarbecovirus positive by RT-qPCR | OZ248189 | NA |
| RshSTT610 | *Rhinolophus shameli* | Phnom Chab Phleurng | 2023-06-17 | 18.36 | 16.45 | Positive | Positive | OZ251056/OZ250977 | BetaCoV-Sarbecovirus | OZ248275 | NA |
| RshSTT612 | *Rhinolophus shameli* | Phnom Chab Phleurng | 2023-06-17 | 40.49 | 38.15 | Negative | Negative | NA | Sarbecovirus positive by RT-qPCR | OZ248276 | NA |
| RshSTT010 | *Rhinolophus shameli* | Phnom Chhngauk | 2023-03-27 | Negative | Negative | Positive | Positive | OZ250982/OZ250903 | AlphaCoV-Decacovirus | OZ248277 | NA |
| RshSTT011 | *Rhinolophus shameli* | Phnom Chhngauk | 2023-03-27 | Negative | Negative | Positive | Positive | OZ250983/OZ250904 | AlphaCoV-Decacovirus | OZ248278 | NA |
| RshSTT012 | *Rhinolophus shameli* | Phnom Chhngauk | 2023-03-27 | Negative | Negative | Positive | Positive | OZ250984/OZ250905 | AlphaCoV-Decacovirus | OZ248279 | NA |
| RshSTT061 | *Rhinolophus shameli* | Phnom Ka Ngoak | 2023-03-28 | Negative | Negative | Positive | Positive | OZ250985/OZ250906 | AlphaCoV-Decacovirus | OZ248280 | NA |
| RshSTT067 | *Rhinolophus shameli* | Phnom Ka Ngoak | 2023-03-28 | Negative | Negative | Negative | Positive | OZ250880 | AlphaCoV-Decacovirus | OZ248190 | NA |
| RshSTT072 | *Rhinolophus shameli* | Phnom Ka Ngoak | 2023-03-28 | Negative | Negative | Positive | Positive | OZ250986/OZ250907 | AlphaCoV-Decacovirus | OZ248281 | NA |
| RshSTT105 | *Rhinolophus shameli* | Phnom Chab Phleurng | 2023-03-29 | Negative | Negative | Positive | Positive | OZ250987/OZ250908 | AlphaCoV-Decacovirus | OZ248282 | NA |
| RshSTT126 | *Rhinolophus shameli* | Phnom Chab Phleurng | 2023-03-29 | Negative | Negative | Negative | Positive | OZ250881 | AlphaCoV-Decacovirus | OZ248283 | NA |
| RshSTT188 | *Rhinolophus shameli* | Phnom Chhngauk | 2023-03-30 | Negative | Negative | Negative | Positive | OZ250882 | AlphaCoV-Rhinacovirus | OZ248284 | NA |
| HpoSTT192 | *Hipposideros pomona* | Phnom Chhngauk | 2023-03-30 | Negative | Negative | Positive | Positive | OZ250988/OZ250909 | AlphaCoV-Decacovirus | OZ248285 | NA |
| RshSTT193 | *Rhinolophus shameli* | Phnom Chhngauk | 2023-03-30 | Negative | Negative | Negative | Positive | OZ250883 | AlphaCoV-Rhinacovirus | OZ248286 | NA |
| RshSTT229 | *Rhinolophus shameli* | Phnom Chhngauk | 2023-05-02 | Negative | Negative | Positive | Positive | OZ250991/OZ250912 | AlphaCoV-Decacovirus | OZ248287 | NA |
| RshSTT276 | *Rhinolophus shameli* | Phnom Chab Phleurng | 2023-05-04 | Negative | Negative | Positive | Positive | OZ250999/OZ250920 | AlphaCoV-Rhinacovirus | OZ248288 | NA |
| RshSTT278 | *Rhinolophus shameli* | Phnom Chab Phleurng | 2023-05-04 | Negative | Negative | Positive | Positive | OZ251000/OZ250921 | AlphaCoV-Decacovirus | OZ248289 | NA |
| RshSTT289 | *Rhinolophus shameli* | Phnom Chab Phleurng | 2023-05-04 | Negative | Negative | Negative | Positive | OZ250886 | AlphaCoV-Decacovirus | OZ248290 | NA |
| RshSTT304 | *Rhinolophus shameli* | Phnom Chab Phleurng | 2023-05-04 | Negative | Negative | Positive | Positive | OZ251009/OZ250930 | AlphaCoV-Decacovirus | OZ248291 | NA |
| RshSTT324 | *Rhinolophus shameli* | Phnom Chhngauk | 2023-05-05 | Negative | Negative | Positive | Positive | OZ251017/OZ250938 | AlphaCoV-Decacovirus | OZ248292 | NA |
| RchSTT332 | *Rhinolophus chaseni* | Phnom Chhngauk | 2023-05-05 | Negative | Negative | Negative | Positive | OZ250891 | AlphaCoV-Decacovirus | OZ248293 | NA |
| HlaSTT383 | *Hipposideros larvatus* | Phnom Chhngauk | 2023-06-12 | Negative | Negative | Positive | Positive | OZ251029/OZ250950 | BetaCoV-Hibecovirus | OZ248294 | NA |
| HlaSTT384 | *Hipposideros larvatus* | Phnom Chhngauk | 2023-06-12 | Negative | Negative | Positive | Positive | OZ251030/OZ250951 | BetaCoV-Hibecovirus | OZ248295 | NA |
| HlaSTT390 | *Hipposideros larvatus* | Phnom Chhngauk | 2023-06-12 | Negative | Negative | Positive | Positive | OZ251031/OZ250952 | BetaCoV-Hibecovirus | OZ248296 | NA |
| HlaSTT416 | *Hipposideros larvatus* | Phnom Ka Ngoak | 2023-06-13 | Negative | Negative | Positive | Positive | OZ251033/OZ250954 | BetaCoV-Hibecovirus | OZ248297 | NA |
| HpoSTT420 | *Hipposideros pomona* | Phnom Ka Ngoak | 2023-06-13 | Negative | Negative | Positive | Positive | OZ251034/OZ250955 | AlphaCoV-Decacovirus | OZ248298 | NA |
| RshSTT433 | *Rhinolophus shameli* | Phnom Ka Ngoak | 2023-06-13 | Negative | Negative | Negative | Positive | OZ250893 | AlphaCoV-Decacovirus | OZ248299 | NA |
| RmiSTT466 | *Rhinolophus microglobosus* | Phnom Chab Phleurng | 2023-06-14 | Negative | Negative | Positive | Positive | OZ251035/OZ250956 | AlphaCoV-Rhinacovirus | OZ248300 | NA |
| RshSTT472 | *Rhinolophus shameli* | Phnom Chab Phleurng | 2023-06-14 | Negative | Negative | Negative | Positive | OZ250895 | AlphaCoV-Decacovirus | OZ248301 | NA |
| HlaSTT489 | *Hipposideros larvatus* | Phnom Chab Phleurng | 2023-06-14 | Negative | Negative | Positive | Positive | OZ251036/OZ250957 | AlphaCoV-Decacovirus & BetaCoV-Hibecovirus | OZ248302 | NA |
| HlaSTT507 | *Hipposideros larvatus* | Phnom Chhngauk | 2023-06-15 | Negative | Negative | Positive | Positive | OZ251040/OZ250961 | AlphaCoV-Decacovirus | OZ248303 | NA |
| HlaSTT508 | *Hipposideros larvatus* | Phnom Chhngauk | 2023-06-15 | Negative | Negative | Positive | Positive | OZ251041/OZ250962 | AlphaCoV-Decacovirus & BetaCoV-Hibecovirus | OZ248304 | NA |
| TmeSTT519 | *Taphozous melanopogon* | Phnom Chhngauk | 2023-06-15 | Negative | Negative | Negative | Positive | OZ250899 | BetaCoV-Hibecovirus | OZ248191 | NA |
| HdiSTT528 | *Hipposideros diadema* | Phnom Chhngauk | 2023-06-15 | Negative | Negative | Positive | Positive | OZ251046/OZ250967 | BetaCoV-Hibecovirus | OZ248305 | NA |
| MspSTT529 | *Megaderma spasma* | Phnom Ka Ngoak | 2023-06-16 | Negative | Negative | Negative | Positive | OZ250901 | BetaCoV-Hibecovirus | OZ248306 | NA |
| RshSTT561 | *Rhinolophus shameli* | Phnom Ka Ngoak | 2023-06-16 | Negative | Negative | Negative | Positive | OZ250902 | AlphaCoV-Decacovirus | OZ248307 | NA |
| HpoSTT572 | *Hipposideros pomona* | Phnom Chab Phleurng | 2023-06-17 | Negative | Negative | Positive | Positive | OZ251051/OZ250972 | AlphaCoV-Decacovirus | OZ248308 | NA |
| HlaSTT605 | *Hipposideros larvatus* | Phnom Chab Phleurng | 2023-06-17 | Negative | Negative | Positive | Positive | OZ251055/OZ250976 | AlphaCoV-Decacovirus | OZ248309 | NA |
| HpoSTT606 | *Hipposideros pomona* | Phnom Chab Phleurng | 2023-06-17 | Negative | Negative | Positive | Negative | OZ250981 | AlphaCoV-Decacovirus | OZ248310 | NA |

NA: Not available

**Supplementary Table S2. List of the 50 SARS-CoV-2 related virus positive bats samples processed for sequencing**

| Sample ID | Bat Genus/Species | Sampling date | Collection site | Sarbecovirus RT-qPCR (19) | | Sequence length (nt) | Virus group |
|---|---|---|---|---|---|---|---|
| | | | | E-gene Ct value | N-gene Ct value | | |
| RshSTT039 | *Rhinolophus shameli* | 2020-08-18 | Phnom Chhngauk | 29.22 | 27.33 | 29735 | 1 |
| RshSTT334 | *Rhinolophus shameli* | 2021-03-15 | Phnom Chhngauk | 24.35 | 23.5 | 29802 | 1 |
| MspSTT571 | *Megaderma Spasma* | 2021-11-30 | Phnom Chab Pleurng | 38.4 | 38.27 | 5767 (partial) | 1 |
| TmeSTT622 | *Taphozous melanopogon* | 2021-12-01 | Phnom Ka Ngoak | 37.99 | 36.38 | 11077 (partial) | 1 |
| RshSTT671 | *Rhinolophus shameli* | 2021-12-02 | Phnom Chhngauk | 21.22 | 19.21 | 29694 | 1 |
| RshSTT779 | *Rhinolophus shameli* | 2021-12-06 | Phnom Ka Ngoak | 23.98 | 22.23 | 29694 | 1 |
| RshSTT783 | *Rhinolophus shameli* | 2021-12-06 | Phnom Ka Ngoak | 34.38 | 33.34 | 20907 (partial) | 1 |
| RshSTT785 | *Rhinolophus shameli* | 2021-12-06 | Phnom Ka Ngoak | 40.63 | 36.77 | 10002 (partial) | 1 |
| RshSTT814 | *Rhinolophus shameli* | 2021-12-06 | Phnom Ka Ngoak | 36.77 | 37.03 | 11415 (partial) | 1 |
| RshSTT219 | *Rhinolophus shameli* | 2023-05-02 | Phnom Chhngauk | 24.29 | 24.02 | 29694 | 1 |
| RshSTT227 | *Rhinolophus shameli* | 2023-05-02 | Phnom Chhngauk | 23.29 | 23.17 | 29694 | 1 |
| RshSTT241 | *Rhinolophus shameli* | 2023-05-03 | Phnom Ka Ngoak | 28.08 | 27.05 | 29694 | 1 |
| RshSTT245 | *Rhinolophus shameli* | 2023-05-03 | Phnom Ka Ngoak | 17.05 | 16.86 | 29694 | 1 |
| RshSTT268 | *Rhinolophus shameli* | 2023-05-04 | Phnom Chab Pleurng | 21.46 | 21.12 | 29694 | 1 |
| RshSTT287 | *Rhinolophus shameli* | 2023-05-04 | Phnom Chab Pleurng | 23.5 | 21.45 | 29694 | 1 |
| RshSTT288 | *Rhinolophus shameli* | 2023-05-04 | Phnom Chab Phleurng | 37.12 | 36.22 | Negative | NA |
| RshSTT307 | *Rhinolophus shameli* | 2023-05-04 | Phnom Chab Phleurng | 36.13 | 33.92 | 1039 (partial) | Partial RdRp |
| RshSTT313 | *Rhinolophus shameli* | 2023-05-05 | Phnom Chhngauk | 36.65 | 36.33 | 1033 (partial) | Partial RdRp |
| RshSTT314 | *Rhinolophus shameli* | 2023-05-05 | Phnom Chhngauk | 20.87 | 18.09 | 29694 | 1 |
| RshSTT317 | *Rhinolophus shameli* | 2023-05-05 | Phnom Chhngauk | 20.15 | 17.34 | 29694 | 1 |
| RshSTT322 | *Rhinolophus shameli* | 2023-05-05 | Phnom Chhngauk | 21.32 | 19.14 | 29694 | 1 |
| RshSTT336 | *Rhinolophus shameli* | 2023-05-05 | Phnom Chhngauk | 19.27 | 17.78 | 29694 | 1 |
| RshSTT344 | *Rhinolophus shameli* | 2023-05-05 | Phnom Chhngauk | 19.05 | 17.33 | 29694 | 1 |
| RshSTT381 | *Rhinolophus shameli* | 2023-06-12 | Phnom Chhngauk | 23.66 | 20.39 | 29694 | 1 |
| RshSTT395 | *Rhinolophus shameli* | 2023-06-12 | Phnom Chhngauk | 24.85 | 23.64 | 29694 | 1 |
| RshSTT419 | *Rhinolophus shameli* | 2023-06-13 | Phnom Ka Ngoak | 19.33 | 18.45 | 29692 | 1 |
| RshSTT494 | *Rhinolophus shameli* | 2023-06-15 | Phnom Chhngauk | 21.08 | 19.68 | 29694 | 1 |
| RshSTT503 | *Rhinolophus shameli* | 2023-06-15 | Phnom Chhngauk | 34.86 | 34.27 | Negative | NA |
| RshSTT515 | *Rhinolophus shameli* | 2023-06-15 | Phnom Chhngauk | 18.65 | 17.22 | 29694 | 1 |
| RshSTT517 | *Rhinolophus shameli* | 2023-06-15 | Phnom Chhngauk | 35.69 | 35.21 | Negative | NA |
| RshSTT523 | *Rhinolophus shameli* | 2023-06-15 | Phnom Chhngauk | 38.18 | 39.38 | Negative | NA |
| RshSTT564 | *Rhinolophus shameli* | 2023-06-16 | Phnom Ka Ngoak | 21.01 | 19.27 | 29694 | 1 |
| RshSTT570 | *Rhinolophus shameli* | 2023-06-17 | Phnom Chab Pleurng | 21.55 | 20.28 | 29821 | 1 |
| RshSTT610 | *Rhinolophus shameli* | 2023-06-17 | Phnom Chab Pleurng | 18.36 | 16.45 | 29694 | 1 |
| RmiSTT243 | *Rhinolophus microglobosus* | 2023-05-03 | Phnom Ka Ngoak | 17.86 | 15.76 | 29694 | 1 |
| RmiSTT249 | *Rhinolophus microglobosus* | 2023-05-03 | Phnom Ka Ngoak | 23.04 | 22.06 | 29694 | 1 |
| RmiSTT252 | *Rhinolophus microglobosus* | 2023-05-03 | Phnom Ka Ngoak | 39.58 | 37.47 | Negative | NA |
| RmiSTT521 | *Rhinolophus microglobosus* | 2023-06-15 | Phnom Chhngauk | 28.29 | 26.39 | 29694 | 1 |
| HlaSTT221 | *Hipposideros larvatus* | 2023-05-02 | Phnom Chhngauk | 39.57 | 38.74 | 8756 (partial) | 1 |
| LlySTT440 | *Lyroderma lyra* | 2023-06-13 | Phnom Ka Ngoak | 37.94 | 36.87 | 16012 (partial) | 1 |
| RchSTT337 | *Rhinolophus chaseni* | 2023-05-05 | Phnom Chhngauk | 37.05 | 38.37 | Negative | NA |
| RchSTT352 | *Rhinolophus chaseni* | 2023-05-05 | Phnom Chhngauk | 38.41 | 37.25 | Negative | NA |
| HarSTT369 | *Hipposideros armiger* | 2023-05-06 | Phnom Ka Ngoak | 38.47 | 38.9 | Negative | NA |
| RacSTT333 | *Rhinolophus acuminatus* | 2023-05-05 | Phnom Chhngauk | 29.07 | 28.72 | 29747 | 2 |
| RacSTT334 | *Rhinolophus acuminatus* | 2023-05-05 | Phnom Chhngauk | 32.12 | 31.42 | 29744 | 2 |
| RacSTT345 | *Rhinolophus acuminatus* | 2023-05-05 | Phnom Chhngauk | 17.68 | 15.52 | 29891 | 2 |
| RacSTT347 | *Rhinolophus acuminatus* | 2023-05-05 | Phnom Chhngauk | 30.51 | 28.78 | 29744 | 2 |
| RacSTT351 | *Rhinolophus acuminatus* | 2023-05-05 | Phnom Chhngauk | 22.13 | 19.66 | 29883 | 2 |
| RpuSTT361 | *Rhinolophus pusillus* | 2023-05-06 | Phnom Ka Ngoak | 31.41 | 33.09 | 29750 | 3 |
| RmaSTT500 | *Rhinolophus malayanus* | 2023-06-15 | Phnom Chhngauk | 27.47 | 28.45 | 29642 | 4 |

NA: Not available. RdRp: RNA-dependent RNA polymerase.

**Supplementary Table S3. List of publicly available sequences used in this study**

| Sequence name | Country | Species | Accession number (GenBank or GISAID) | Collection date |
|---|---|---|---|---|
| BANAL-20-103 | Laos | Bat-*R_pusillus* | MZ937001 | 2020-07-07 |
| BANAL-20-116 | Laos | Bat-*R_malayanus* | MZ937002 | 2020-07-07 |
| BANAL-20-236 | Laos | Bat-*R_marshalli* | MZ937003 | 2020-07-10 |
| BANAL-20-247 | Laos | Bat-*R_malayanus* | MZ937004 | 2020-07-10 |
| BANAL-20-52 | Laos | Bat-*R_malayanus* | MZ937000 | 2020-07-05 |
| CoVZC45 | China-Zhoushan-Dinghai | Bat-*R_sinicus* | MG772933 | 2017-02 |
| CoVZXC21 | China-Zhoushan-Dinghai | Bat-*R_sinicus* | MG772934 | 2015-07 |
| HN2021A | China-Hunan | Bat-*R_pusillus* | OK017803 | 2021-02 |
| HN2021B | China-Hunan | Bat-*R_pusillus* | OK017804 | 2021-02 |
| HN2021G | China-Hunan | Bat-*R_pusillus* | OK017805 | 2021-03 |
| PrC31 | China-Yunnan | Bat-*R_pusillus* | MW703458 | 2018-08 |
| Ra22QT106 | Vietnam-Quang Tri | Bat-*R_affinis* | OR233322 | 2022-11-03 |
| Ra22QT135 | Vietnam-Quang Tri | Bat-*R_affinis* | OR233323 | 2022-11-04 |
| Ra22QT137 | Vietnam-Quang Tri | Bat-*R_affinis* | OR233328 | 2022-11-04 |
| Ra22QT77 | Vietnam-Quang Tri | Bat-*R_affinis* | OR233324 | 2022-11-03 |
| RacCS203 | Thailand-Chachoengsao | Bat-*R_acuminatus* | MW251308 | 2020-06-19 |
| RaTG13 | China-Yunnan | Bat-*R_affinis* | MN996532 | 2013-07-24 |
| RmYN02 | China-Yunnan | Bat-*R_malayanus* | EPI_ISL_412977 | 2019-06-25 |
| Rp_JCC9 | China-Yunnan | Bat-*R_blythi* | OK287355 | 2020-03 |
| Rp22DB159 | Vietnam-Dien Bien | Bat-*R_pusillus* | OR233302 | 2022-06-26 |
| RpYN06 | China-Yunnan | Bat-*R_pusillus* | MZ081381 | 2020-05-25 |
| Wuhan-Hu-1 | China-Wuhan | Human | NC_045512 | 2019-12 |
| YN2021 | China-Yunnan | Bat-*R_pusillus* | OK017806 | 2021-04 |
| R24 | China-Yunnan | Bat-*R_marshalli* | OP963576 | 2018 |
| G1 | China-Guangdong | Pangolin | EPI_ISL_410721 | 2019 |
| MP789 | China-Guangdong | Pangolin | MT121216 | 2019-03-29 |
| P1E | China-Guangxi | Pangolin | MT040334 | 2017 |
| P2V | China-Guangxi | Pangolin | MT072864 | 2018 |
| P4L | China-Guangxi | Pangolin | MT040333 | 2017 |
| P5E | China-Guangxi | Pangolin | MT040336 | 2017 |
| P5L | China-Guangxi | Pangolin | MT040335 | 2017 |
| P79-9 | China-Guangdong | Pangolin | OQ297708 | 2019-03 |
| RshSTT182 | Cambodia-Steung Treng | Bat-*R_shameli* | EPI_ISL_852604 | 2010-12-06 |
| RshSTT200 | Cambodia-Steung Treng | Bat-*R_shameli* | EPI_ISL_852605 | 2010-12-06 |

*R= Rhinolophus*

**Supplementary Table S4. Evaluation of temporal signal and clock models comparisons**

| Dataset | Clock model | Log marginal likelihood with sampling dates | Log marginal likelihood without sampling dates |
|---|---|---|---|
| R1 | SC | -14918 | -14970 |
| | UCLN | -14894 | -14918 |
| R2 | SC | -9854 | -9893 |
| | UCLN | -9824 | -9843 |
| R3 | SC | -13024 | -13052 |
| | UCLN | -12955 | -12977 |
| R4 | SC | -19341 | -19377 |
| | UCLN | -19043 | -19061 |
| R5 | SC | -6512 | -6575 |
| | UCLN | -6483 | -6517 |
| R6 | SC | -10615 | -10664 |
| | UCLN | -10588 | -10606 |
| R7 | SC | -6566 | -6594 |
| | UCLN | -6565 | -6596 |
| R8 | SC | -9051 | -9097 |
| | UCLN | -9025 | -9052 |
| R9 | SC | -10172 | -10176 |
| | UCLN | -10152 | -10171 |
| R10 | SC | -8577 | -8623 |
| | UCLN | -8540 | -8550 |
| R11 | SC | -6229 | -6273 |
| | UCLN | -6205 | -6234 |
| R12 | SC | -13997 | -14022 |
| | UCLN | -13827 | -13846 |
| R13 | SC | -6366 | -6382 |
| | UCLN | -6349 | -6362 |
| NRA | SC | -95382 | -95468 |
| | UCLN | -95364 | -95388 |
| Non recombinant region group 1 | SC | -43407 | -43493 |
| | UCLN | -43403 | -43424 |
| Non recombinant region group 1a | SC | -40830 | -40889 |
| | UCLN | -40830 | -40866 |

We tested two molecular clock models. SC: Strict clock, UCLN: uncorrelated relaxed clock with lognormal distribution.

Log-marginal likelihood estimates using the generalized stepping stone sampling (GSS) model selection approach are presented for dataset using the sample dates, and the same dataset analyzed without dates, resulting in positive Bayes factors and indicating that there was temporal signal in the data.

### Supplementary Table S5. Coalescent tree priors comparisons

| Dataset | | Coalescent tree prior Log marginal likelihood | | | | |
|---|---|---|---|---|---|---|
| | | Constant size | Exponential growth | Bayesian skygrid | Bayesian skyline | Bayesian skyride |
| R1 | PS | -14851 | -14848 | -14810 | **-14807** | -14842 |
| | SS | -14852 | -14849 | -14811 | **-14807** | -14842 |
| R2 | PS | -9810 | -9801 | -9762 | **-9760** | -9800 |
| | SS | -9811 | -9801 | -9763 | **-9761** | -9800 |
| R3 | PS | -12910 | -12909 | -12875 | **-12872** | -12904 |
| | SS | -12911 | -12910 | -12877 | **-12873** | -12904 |
| R4 | PS | -18925 | -18920 | -18884 | **-18880** | -18908 |
| | SS | -18926 | -18921 | -18885 | **-18881** | -18908 |
| R5 | PS | -6464 | -6461 | -6443 | **-6440** | -6468 |
| | SS | -6465 | -6462 | -6443 | **-6441** | -6468 |
| R6 | PS | -10545 | -10544 | -10502 | **-10493** | -10541 |
| | SS | -10546 | -10545 | -10503 | **-10494** | -10541 |
| R7 | PS | -6536 | -6531 | -6492 | **-6487** | -6537 |
| | SS | -6537 | -6532 | -6493 | **-6488** | -6537 |
| R8 | PS | -9007 | -9004 | -8969 | **-8963** | -9006 |
| | SS | -9008 | -9005 | -8969 | **-8964** | -9006 |
| R9 | PS | -10140 | -10139 | -10094 | **-10089** | -10141 |
| | SS | -10141 | -10140 | -10094 | **-10091** | -10141 |
| R10 | PS | -8525 | -8524 | -8492 | **-8489** | -8528 |
| | SS | -8526 | -8525 | -8493 | **-8490** | -8528 |
| R11 | PS | -6196 | -6193 | **-6158** | -6161 | -6191 |
| | SS | -6197 | -6193 | **-6158** | -6161 | -6191 |
| R12 | PS | -13774 | -13772 | -13732 | **-13726** | -13752 |
| | SS | -13775 | -13772 | -13733 | **-13726** | -13752 |
| R13 | PS | -6340 | -6337 | -6305 | **-6304** | -6329 |
| | SS | -6340 | -6337 | -6305 | **-6304** | -6329 |
| NRA | PS | -94937 | -94930 | -94893 | **-94881** | -94931 |
| | SS | -94938 | -94931 | -94895 | **-94880** | -94930 |
| Non recombinant region group 1 | PS | **-43360** | **-43360** | **-43360** | **-43360** | -43366 |
| | SS | **-43360** | **-43360** | **-43360** | **-43360** | -43366 |
| Non recombinant region group 1a | PS | -40821 | -40822 | **-40820** | -40826 | -40822 |
| | SS | -40821 | -40822 | **-40820** | -40826 | -40822 |

The log marginal likelihood values were estimated using both path sampling (PS) and stepping-stone sampling (SS), and the values highlighted in bold indicate the best supported model from log Bayes factor comparisons.
